# Structural differences in translation initiation between pathogenic trypanosomatids and their mammalian hosts

**DOI:** 10.1101/806141

**Authors:** Anthony Bochler, Jailson Brito Querido, Terezie Prilepskaja, Heddy Soufari, Angelita Simonetti, Mayara Lucia Del Cistia, Lauriane Kuhn, Aline Rimoldi Ribeiro, Leoš Shivaya Valášek, Yaser Hashem

## Abstract

Canonical mRNA translation in eukaryotes begins with the formation of the 43S pre-initiation complex (PIC). Its assembly requires the binding of several eukaryotic initiation factors (eIF 1, 1A, 2, 3 and 5), Met-tRNA_i_^Met^ and the small ribosomal subunit (40S). Compared to their mammalian hosts, trypanosomatids present significant structural differences in their 40S suggesting substantial variability in translation initiation. Here we determined the structure of the 43S PIC from *Trypanosoma cruzi*, the parasite causing the Chagas disease. Our structure shows numerous specific features, such as the variant eIF3 structure and its unique interactions with the large rRNA ESs 9^S^, 7^S^ and 6^S^, and the association of a kinetoplastid-specific ~245 kDa DDX60-like helicase. It also revealed the so-far-elusive 40S-binding site of the eIF5 C-terminal domain and the structures of key terminal tails of several conserved eIFs underlying their activities within the PIC. Our results are corroborated by GST-pulldown assays in both human and *T. cruzi* and mass-spectrometry data.

## INTRODUCTION

The first critical initiation step in eukaryotes is the assembly of the 43S PIC comprising the 40S, the eIF2•GTP•Met-tRNA_i_^Met^ ternary complex, and eIFs 1, 1A, 3 and 5 (Valásek, 2012; Hinnebusch, 2017). It is followed by the recruitment of the mRNA promoted by the mRNA cap-binding complex comprising eIF4A, 4B and 4F (Guca et al., 2018; Hashem and Frank, 2018), forming the 48S PIC. The 48S PIC then scans the 5’ untranslated region (UTR) of mRNA in the 5’ to 3’ direction till a start codon is encountered, upon which the majority of eIFs sequentially disassemble from the 40S and the resulting 48S initiation complex (48S IC) joins the large ribosomal subunit (60S) to form an elongation-competent 80S ribosome.

Kinetoplastids is a group of flagellated unicellular eukaryotic parasites that have a complex life cycle. They spend part of their life cycle in the insect guts before being transmitted to the mammalian host upon biting. Common kinetoplastids include human pathogens such as *Trypanosoma cruzi, Trypanosoma brucei* and *Leishmania spp*., etiologic agents of Chagas disease, African sleeping sickness and leishmaniasis, respectively. However, most of the related public health measures are mainly preventative and therapeutic strategies are extremely limited and often highly toxic. Since kinetoplastids have diverged early from other eukaryotes, their mRNA translational machineries developed unique molecular features unseen in other eukaryotic species. For instance, their 40S contains a kinetoplastid-specific ribosomal protein (KSRP) (Brito Querido et al., 2017) and unusually oversized ribosomal RNA (rRNA) expansion segments (ES^S^) (Hashem et al., 2013a). Since these unique features may play specific roles in kinetoplastidian mRNA translation, they provide potential specific drug targets.

It was proposed that two particularly oversized expansion segments, ES6^S^ and ES7^S^ located near the mRNA exit channel on the kinatoplastidian 40S, may contribute to modulating translation initiation in kinetoplastids by interacting with the structural core of the eukaryotic eIF3, specifically via its subunits a and c (Hashem et al., 2013b). eIF3 is the most complex eIF promoting not only nearly all initiation steps, but also translation termination, stop codon readthrough and ribosomal recycling (Valášek et al., 2017). Among its initiation roles, eIF3 critically contributes to the assembly of the 43S PIC through a multitude of contacts that it makes with other eIFs, ensuring their recruitment to the 40S (Valášek et al., 2017). Mammalian eIF3 comprises twelve subunits (eIF3a–m; excluding j), eight of which form the PCI/MPN octameric structural core (eIF3a, c, e, f, h, k, l and m) (des Georges et al., 2015; Sun et al., 2011; Wagner et al., 2014, 2016). Interestingly, unlike their mammalian hosts, kinetoplastids do not encode the eIF3m subunit (Li et al., 2017; Rezende et al., 2014; Meleppattu et al., 2015) co-forming the octameric core in all known “12-subunit” species, strongly suggesting that the structure of their eIF3 core differs from that of mammals.

The 43S PIC assembly is also enhanced by the C-terminal domain (CTD) of eIF5 (Asano et al., 2001). Indeed, biochemical and genetics studies revealed that the eIF5-CTD possesses specific motifs interacting with several eIFs, such as the N-terminal tail (NTT) of the β subunit of eIF2(Asano et al., 1999; Das et al., 1997). However, the molecular details underlying the eIF5-CTD critical assembly role remain elusive, and – in contrast to the eIF5-NTD (Llácer et al., 2018) – so are the structural details of its binding site within the 43S PIC (Zeman et al., 2019). Importantly, structures of terminal tails of several essential eIFs in most of the available cryo-EM reconstructions are also lacking, mainly due to their intrinsic flexibility. Among them stand out the terminal tails of the c and d subunits of eIF3, eIF2β, eIF1 and eIF1A, all critically involved in scanning and AUG recognition.

Here, we solved the structure of the 43S PIC from *Trypanosoma cruzi* at 3.33Å and unraveled various new aspects of this complex, some of which are specific to trypanosomatids and others common to eukaryotes. Our structures thus allow us to 1) pin point essential, specific-features of trypanosomatids that could represent potential drug targets, and 2) expand our understanding of the interaction network between several eIFs within the 43S PIC underlying molecular mechanism of its assembly, as well as of their roles in scanning for start codon recognition.

## RESULTS AND DISCUSSION

### Composition of the 43S PIC in trypanosomatids

We purified endogenous pre-initiation complexes from two different species, *Trypanosoma cruzi* and *Leishmania tarentolae* by stalling the 43S complexes with GMP-PNP, a non-hydrolysable analog of GTP, as previously described (Simonetti et al., 2016; Simonetti & Guca et al. 2020). The proteomic analysis comparison between the stalled *versus* untreated complexes from *T. cruzi* indicated an obvious enrichment in canonical eIFs and ABCE1, as expected (see methods, Fig. 1A-B and Supplementary Fig. 1). Surprisingly, we also identified an orthologue of the human DEAD-box RNA helicase DDX60 (Fig. 1B, Supplementary Fig. 1). A similar repertoire of eIFs can also be found in the 43S PIC from *L. tarentolae* (Fig. 1B, Supplementary Fig. 2). Besides initiation factors, several other proteins contaminating the 43S PIC can be found in *T. cruzi* and *L. tarentolae* samples without any apparent link to the translation process. Noteworthy, to date and to the best of our knowledge, DDX60 has never been co-purified with any PICs from any other studied eukaryote. Interestingly, while DDX60 is non-essential in mammals (Miyashita et al., 2011; Oshiumi et al., 2015), it is required for the cell fitness in kinetoplastids and trypanosomatides (Alsford et al., 2011), indicating that it could play a specific role in translation initiation in these parasites. It is not known whether or not it is essential in yeast.

**Figure 1.**
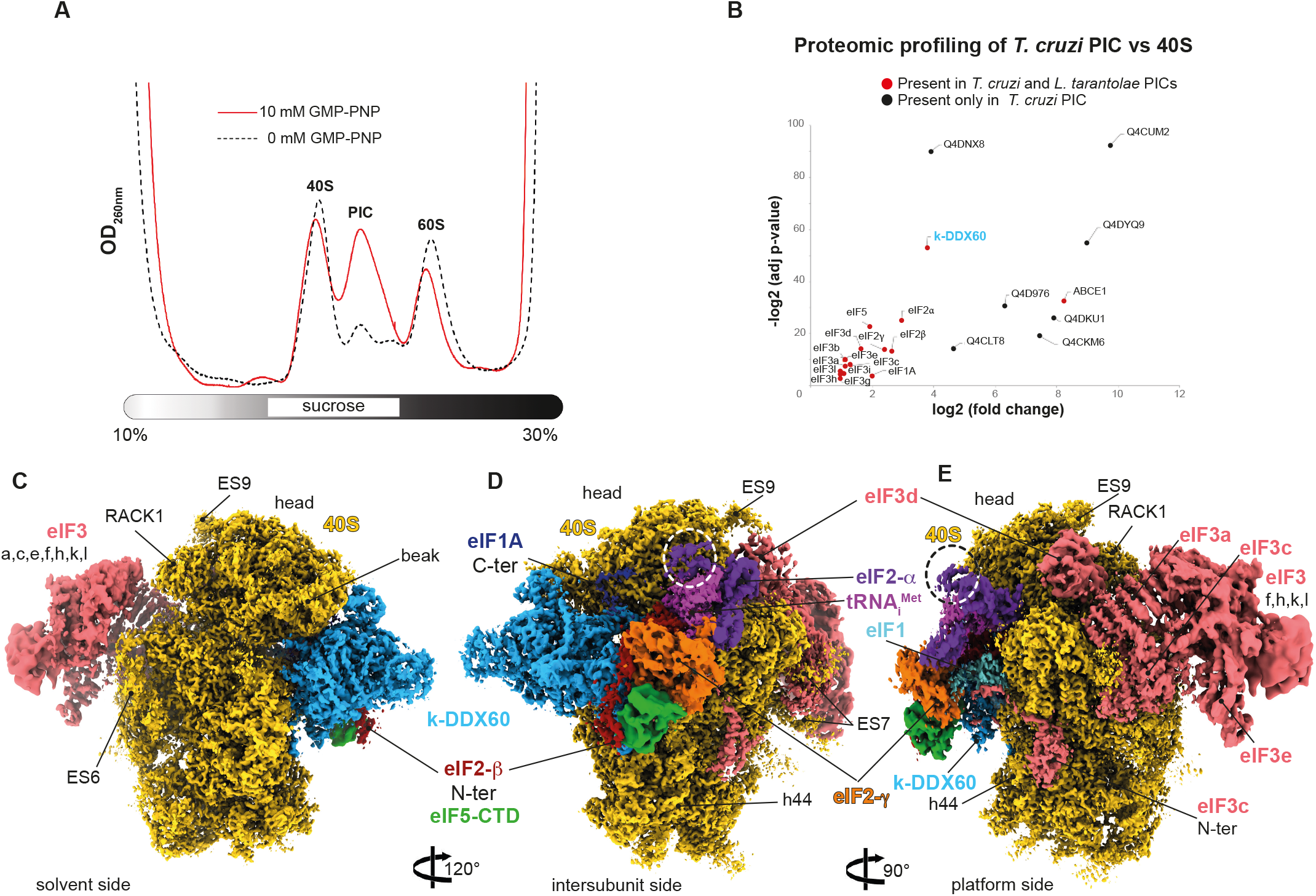
Composition and cryo-EM structure of the T. cruzi 43S PIC. (**A**) The effect of the GMP-PNP treatment on the 43S PIC stabilization in the *T.cruzi* lysate assessed by UV absorbance profile analyses (**B**) Proteomic profiling of the endogenous pre-initiation complex in comparison with native 40Ss purified from the *T. cruzi* cell lysate (see methods for the validation). (**C**) The overall structure of the *T. cruzi* 43S PIC shown from the intersubunit side. The initiation factors are colored variably. (**D**) The 43S PIC reconstruction focused on the solvent side. Extra density of eIF2α corresponding to the kinetoplastidian specific N-terminal insertion is encircled by a dashed line. (**E**) The 43S PIC reconstruction focused on eIF3 and the 40S platform. Different segments are were colored according to their average local resolutions.

### The cryo-EM structure of the 43S PIC from T. cruzi

We next employed cryo-electron microscopy (cryo-EM) to determine the structure of the *T. cruzi* 43S PIC to an overall resolution of 3.33Å, after image processing and extensive particle sorting (Supplementary Fig. 3 and 4). Our reconstruction reveals the so-called “scanning-conducive conformation” of the 43S PIC, in which the head of the 40S is tilted upwards to open up the mRNA channel for the subsequent mRNA loading (Llácer et al., 2015; des Georges et al., 2015; Hashem et al., 2013). Thanks to the conservation of structures and binding sites of most of the identified initiation factors, we were able to segment the map accordingly, thus yielding density segments corresponding to the 40S, eIF1, eIF1A, eIF2α, eIF2β, eIF2g, Met-tRNA_i_^Met^ and the eIF3 structural core (Fig. 1C-E). Importantly, we could also identify the entire density corresponding to the N-terminal tail of the eIF3d subunit, implicated in the mRNA-specific translational control (Lee et al., 2015, 2016) (see below).

Furthermore, we observed an unassigned density contacting eIF2g that has not been seen previously in any equivalent complexes. Since rigid body fitting of the crystal structure of the eIF5-CTD (Wei et al., 2006) showed a close agreement with this unassigned density and previous biochemical and genetics findings suggested a close co-operation between eIF5 and eIF2 on the ribosome (Asano et al., 1999; Luna et al., 2012; Singh et al., 2004, 2012), we assigned this density to the eIF5-CTD (Fig. 1C-E). Because the eIF5-CTD is known to interact with the eIF2β-NTT in both yeasts and mammals (Asano et al., 1999; Das et al., 1997; Das et al., 2000), we could also for assign part of the eIF2β-NTT to its corresponding density (Fig. 1D) (see below). It is important to highlight that it was possible to assign the above-mentioned densities to eIF5-CTD thanks to its general conservation among eukaryotes.

As discussed in detail below, beyond these evolutionary conserved features of the 43S PIC in eukaryotes, our cryo-EM reconstruction also identified several trypanosomatide and kinetoplastid-specific peculiarities. For instance, the kinetoplastidian eIF2α contains a specific N-terminal domain insertion of unknown function (Supplementary Fig. 5A), and, indeed, an extra density on the eIF2α subunit can be observed (Fig. 1D-E, dashed circle). We also revealed a large density at the 40S interface, in the vicinity of the mRNA channel entrance (Fig. 1C-D), unseen in any of the previous mammalian and yeast 43S PIC reconstructions. Taken into account our proteomic analysis (Fig. 1B and Supplementary Fig. 1 and 2), the size of this additional density and, above all, its high-resolution features, we were able to assign it unambiguously to the kinetoplastidian DDX60 (k-DDX60) helicase. These same k-DDX60 and eIF2α-NTT densities are also present in the *L. tarentolae* 43S PIC reconstruction (Supplementary Fig. 6).

Based on the cryo-EM reconstruction of the *T. cruzi* 43S PIC and the conservation of the initiation factors, a near-complete atomic model was generated (see Methods). Our structure reveals a wealth of new interactions (Table S1, Supplementary Fig. 7 and 8)

### The elF5 C-terminal domain (CTD) in the context of the 43S PIC

Importantly, detailed inspection of our structure allowed us to determine the eIF5-CTD binding site on the 43S PIC. It sits in a pocket formed by the eIF2β-NTT and eIF2g (Fig. 2A-D). It was proposed that the three conserved poly-lysine stretches (dubbed “K-boxes”) within the eIF2β-NTD mediate the eIF2 interaction with the eIF5-CTD (Asano et al., 1999; Das et al., 1997). Interestingly, the K1 and K2 –boxes are conserved in their basic charge character but replaced by R-rich stretches in kinetoplastids (Supplementary Fig. 9). However, as our structure of eIF2β-NTT is only partial, we cannot validate their involvement in the interaction with eIF5. In contrast, the K3-box is not conserved in sequence among kinestoplastids (Supplementary Fig. 9), it is replaced by a Q-rich motif, yet its position and orientation towards its binding partner in the eIF5-CTD is conserved. Additionally, our structure shows numerous other contacts between hydrophobic and charged residues on each side (residues L120, N118, L123, L120, L142, K125 and V132 of eIF2β contact A262 R265, V325, V329, I332, Q364 and W372 of eIF5, respectively) (Fig. 2A, 2B, Supplementary Fig. 7I and Table S1 for details). Since the eIF5 residues 320 through 373 correspond to the conserved and essential segment (known as the bipartite motif – AA (acidic/aromatic)-box1 and 2; Fig. 2 A and B, table S1), which was previously implicated in mediating the eIF5-CTD – eIF2β-NTT contact in both yeast and mammals (Asano et al., 1999, 2001; Das et al., 2000), our structure not only provides critical structural evidence supporting earlier biochemical and genetics analysis, it also clearly indicates that the molecular determinants of the eIF5-CTD–eIF2β-NTT contact are conserved. Therefore, we suggest that the eIF5-CTD occupies the same position also in yeast and mammals.

**Figure 2.**
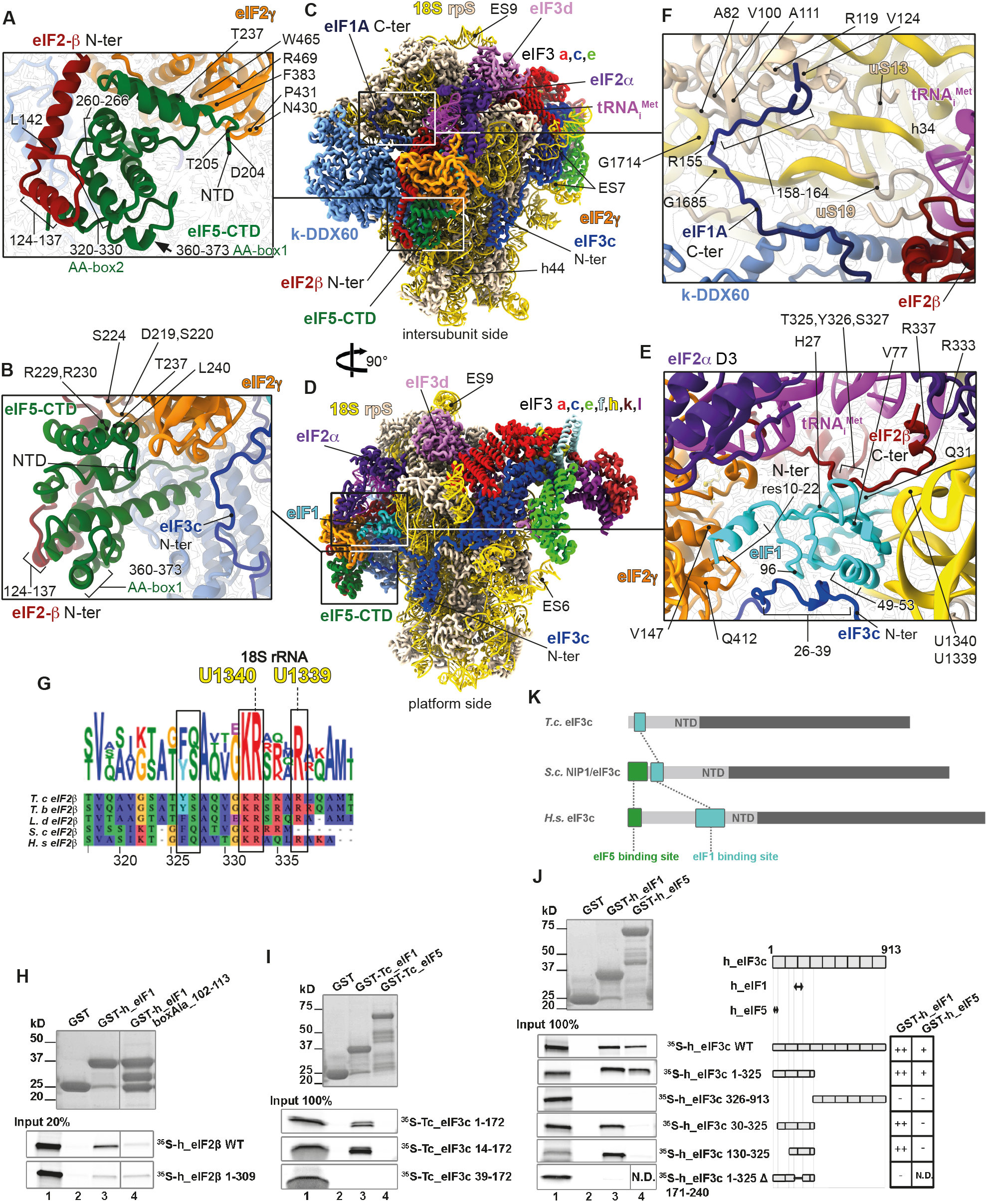
Atomic model of the 43S PIC showing the interaction network of various eIFs. (**A**) Close-up view of an atomic model of the eIF5-CTD (in green), the eIF2β-NTT (in cherry red) and eIF2γ (in orange) shown from the intersubunit side. (**B**) Close-up view of the eIF5-CTD (in green) and its interaction with eIF2 from the platform side. (**C**) The overall view of atomic model of the 43S PIC from the intersubunit and **(D)** the platform side. (**E**) Close-up view of the P-site, showing eIF1 (in cyan) and its biding partners the eIF2β-CTT (in cherry red) and the eIF3c-NTD (in blue). (**F**) Close-up view of the eIF1A-CTT and its interactions with h34, uS13 and uS19. (**G**) Polypeptide sequence alignment of the eIF2β-CTT, highlighting residues involved in the interaction with 18S rRNA and eIF1; *T. cruzi, T. brucei, L. donovani, S. cerevisiae* and *H. sapiens*. Residue numbreding from *H. sapiens* was used (**H**) *In vitro* protein-protein binding analysis of the interaction between human eIF2β and GST-eIF1. (**I**) Binding analysis between the *T. cruzi* eIF3c-NTD and GST-eIF1 and GST-eIF5. (**J**) Binding analysis between human eIF3c-NTD and GST-eIF1 and GST-eIF5. (**K**) Schematics illustrating the differences in the localization of eIF1 (turquoise box) and eIF5 (green box) binding sites within the N-terminal segment of eIF3c in *Trypanosoma cruzi (T.c.), Saccharomyces cerevisiae (S.c.), and Homo sapiens (H.s*.).

Our structure also provides important molecular insight into the eIF5-CTD interaction with the eIF2g domain I (G-domain), where Arg229, Arg230 and R273 of eIF5 contact the G-domain’s Asp219, Ser224 and Ser220, respectively (Fig. 2 A and B, Supplementary Fig. 7 J and K, Table S1 for details). The eIF5-CTD also binds domain III, where Asp204, Thr205, Thr237 and Leu240 of eIF5 interact with domain III’s Pro431, ArgR469-Asn430, Trp465 and Phe383. Noteworthy, the eIF5-CTD shares a common topology with the CTD of the ε subunit of the nucleotide exchange factor eIF2B (Asano et al., 1999); they both fold into a W2-type HEAT domain (Wei et al., 2006) mediating contacts of both factors with the eIF2β-NTT and eIF2g (Alone et al., 2006). Based on our structure, the arrangement of the eIF5-CTD HEAT domain binding site on eIF2g in the context of the 43S PIC is similar to that of the eIF2Bε-CTD HEAT domain in the context of the recently solved eIF2-eIF2B complex (Kenner et al., 2019; Kashiwagi et al., 2019).

Taken together, the eIF5-CTD interaction network revealed here indicates that the interaction between eIF5-CTD and eIF2g could in principle induce a subtle conformational change in its G-domain, allowing the eIF5-NTD (a GTPase activating domain of eIF5) to gain access to the GTP-biding pocket to promote reversible GTP hydrolysis on eIF2 during scanning, as demonstrated earlier (Algire et al., 2005).

### Extensive interaction network of eIF1 in the context of the 43S PIC

After the GTP hydrolysis by eIF2g, the release of the inorganic phosphate (Pi) is prevented by eIF1 until an AUG start codon is recognized by the anticodon of Met-tRNA_i_^Met^ leading to the full accommodation of TC in the decoding pocket (Hinnebusch, 2017; Algire et al., 2005) and eIF1 replacement by the eIF5-NTD. Because the access to the GTP-binding pocket on eIF2g is in part protected by the zinc-binding domain (ZBD) of the eIF2β-CTD (Llácer et al., 2015; Stolboushkina et al., 2008), and biochemical and genetic analysis in yeast indicated that the eIF1 interactions with eIF2β and the NTD of the c subunit of eIF3 play a critical a role in anchoring of eIF1 within the 48S PIC (Thakur et al., 2019; Obayashi et al., 2017; Karásková et al., 2012; Valásek et al., 2004), for our complex understanding of the AUG recognition process it is necessary to investigate how eIF1 coordinates the release of Pi with the latter factors on the molecular level.

In accord with earlier biochemical experiments, our structure reveals that the conserved eIF2β-C terminal tail (eIF2β-CTT), together with the eIF3c-NTD, does anchor eIF1 within the 43S PIC (Fig. 2E). In particular, the eIF2β-CTT extends toward the P-site, where its Thr325, Tyr326 and Ser327 residues interact with eIF1 mainly through His27, Val77 and Gln31, all conserved in character (Fig. 2G, Supplementary Fig. 7A and Table S1). The eIF2β-CTT also interacts with h24 of the 18S rRNA (Arg 333 and 337 with nucleotides U1340, G1342 and U1339) (Fig. 2 E and G, Supplementary Fig. 7H and Table S1). In addition, the eIF2β-binding platform of eIF1 also consists of R29, Q32 and Q43 (Supplementary Fig. 7A and Table S1 for details), as well as of the tip of the eIF1 C-terminus (residues 105-108). Based on these findings, we examined binding of human eIF2β with eIF1 fused to GST moiety using the GST pull down assay and revealed that the interaction between the CTTs of eIF2β (residues 310 – 333) and eIF1 is also conserved in mammals and requires the extreme C-terminus (Fig. 2H, Supplementary Fig. 10A).

The protein sequence composition of the N-terminal domain of eIF3c can vary across species (Supplementary Fig. 11A). It begins with a few conserved hydrophobic residues followed by negatively charged SD/SE repeats in all, including in kinetoplastids. Interestingly, budding yeast *S. cerevisiae* contains an insertion of approximately 40 residues between the latter two groups. The minimal eIF5-CTD binding site within the yeast eIF3c-NTD was identified to fall into the region of the first 45 residues, including part of this insertion but completely excluding the SD/SE repeats (Karásková et al., 2012). These regions are then followed by the segment that was shown to represent the core eIF1-binding segment in yeast (residues 59-87) (Obayashi et al., 2017; Karásková et al., 2012). The downstream sequence in mammals features a specific insertion (residues 167-238), consisting of two highly acidic regions separated by a mostly positively charged/hydrophobic region (Supplementary Fig. 11A). Strikingly, the first part of this mammalian-specific insertion displays a significant sequence similarity with the *S.cerevisiae* core eIF1-binding region; in particular the yeast residues 51-92 show ~36% identity with human residues 173-213 (Supplementary Fig. 11A).

Based on our structure, the contact between the *T.c*. eIF3c-NTD and eIF1 involves Arg26 through Thr39 of eIF3c, and Asn96 and Leu49 through Arg53 of eIF1 (Fig. 2E, Supplementary Fig. 7C and Table S1). In accord, *T.c*. eIF1 fused to GST moiety interacted specifically with the eIF3c-NTD also *in vitro* (the first 14 residues of eIF3c are not required, whereas the following residues up to position 39 are) (Fig. 2I). This interacting region following the extreme N-terminal hydrophobic residues and negatively charged SD/SE repeats nicely correlates with the eIF1 binding region of the *S.cerevisiae* eIF3c-NTD specified above (Obayashi et al., 2017; Karásková et al., 2012) (Supplementary Fig. 11A).

As for the eIF5-CTD– eIF3c-NTD contact, which was so far also determined only in yeast *S. cerevisiae* (Obayashi et al., 2017; Karásková et al., 2012; Valásek et al., 2004; Phan et al., 1998), given the evolutionary conservation of this extreme N-terminal region, one would expect it to be conserved among all eukaryotes too. Therefore, it was rather surprising not to detect any binding between the *T. cruzi* eIF3c-NTD and eIF5 under any *in vitro* experimental set-up at any condition that we examined exhaustively (Fig. 2I and Supplementary Fig. S11 B and C). This is consistent with our structure (Figure 2B), where despite the observable proximity between the eIF3c-NTD and eIF5-CTD, these two domains remain out of the intermolecular interactions range, and for which we detected no structural evidence. Even though we cannot rule out that they may come in contact in the preinitiation complexes only in some stages of the initiation pathway that we did not capture, we tend to think that these results point to a specific evolutionary shift in kinetoplastidian initiation pathway, as will be discussed below.

This unexpected finding prompted us to investigate the conservation of the eIF3c-NTD interactions in higher eukaryotes. Therefore, we fused human eIF1 and eIF5 to GST and tested the resulting fusion proteins against various truncations of the eIF3c-NTD (Fig. 2J). In accord with the yeast data (Obayashi et al., 2017; Karásková et al., 2012) but in contrast to *T. cruzi* (Fig. 2I), the extreme N-terminal group of conserved hydrophobic residues of human eIF3c-NTD interacted strongly with eIF5.

Taken into account the peculiarity of the human eIF3c-NTD featuring the aforementioned insertion (residues 167-238), indicating that the eIF1-binding site appears to be located more towards the C-terminal part of the eIF3c-NTD, we first deleted the first 130 resides and, indeed, showed that the eIF3c-NTD segment spanning residues 130 through 325 fully preserved its affinity towards eIF1 (Fig. 2J). Conversely, internal deletion of residues 171 through 240 from the human eIF3c-NTD construct resulted in a complete loss of binding (Fig. 2J). Thus, the core eIF1-binding site in the human eIF3c-NTD seems to fall into the first part of this mammalian-specific insertion, displaying a significant sequence similarity with the *S.cerevisiae* core eIF1-binding region (Supplementary Fig. S11A), as described above.

Taken together, these findings suggest that despite the undisputable importance of the eIF3c-NTD during the initiation and start-codon recognition, this region has undergone rather dramatic topological, as well as sequential restructuring during the course of evolution. 1) The eIF1 binding site preserved its key sequence determinants but moved further downstream in the course of evolution of higher eukaryotes (Fig. 2K). In contrast, 2) the eIF5 binding site remained conserved not only in its sequence but also in its placement at the extreme N-terminal tip of eIF3c across species, however, in kinetoplastids it most probably lost its purpose. It remains to be seen what molecular consequences of this evolutionary shift are in kinetoplastids and whether or not these two molecules come into a functional contact within the PICs.

Besides the eIF1-CTT binding coordinates, our structure also reveals that the N-terminal tail of eIF1 (residues 10 to 22) forms an α-helix that interacts with domains I and III of eIF2γ (Val85, Val147, Gln412 and Asn459, Fig. 2E, Supplementary Fig. 7B and Table S1), very close to the GTP binding pocket. We propose that these contacts could underlie the role of eIF1 in releasing the Pi by inducing a subtle conformational change in the GTP binding pocket upon sensing the recognition of the start codon through its apical β-hairpin loop at the P-site.

Finally, even though eIF1A appears to interact with eIF1 in a canonical fashion seen in other eukaryotes, it shows that the eIF1A-CTT extends towards the head of the 40S, where it interacts with the rRNA (Arg155 with G1685) (Fig. 2F, Supplementary Fig. 7D and Table S1) and ribosomal proteins uS19 (residues Val158 with Val100, Ala82 and Ala111, Supplementary Fig. 7F and Table S1) and uS13 (residues Asp162 and Leu164 with Arg119 and Val124, respectively; Supplementary Fig. 7G and Table S1), corroborating findings from a previous hydroxyl-radical probing study (Yu et al., 2009). Moreover, previously uncharacterized interactions between eIF1A and eIF2β are observed in T.c. between hydrophobic residues (Tyr133 through Phe135 on eIF1A and Leu282 through Tyr279 on eIF2β, Supplementary Fig. 7E and Table S1)

### The specific features and binding site of eIF3 in trypanosomatids

Strikingly, as seen in Figure 3A-D, the unusually large trypanosomatids-specific ES^S^ are involved in translation initiation by acting as docking platforms for different subunits of eIF3. Similarly to other eukaryotes reported so far, the eIF3 core binds to the 40S through its a and c subunits (Fig. 3C-D). However, unlike in other known eukaryotes, the large ES7^S^ acts as the main docking point for the eIF3 structural core (Supplementary Fig. 12 and 13A). In particular, the eIF3c is tweezed between ES7^S^-helix A (ES7^S^-hA) and ES7^S^-hB forming a large, kinetoplastid-specific binding site, involving residues Gln204, Lys207, Arg215, Arg232, Arg243, Gln329 and Arg331 and ES7^S^ nucleotides U1526, A1525 and U1523, U1476, U1526, G1438 and U1439, respectively (Fig. 3D, Supplementary Fig. 7 M and N, Table S1). The local resolution of our complex allowed us to assign the identity of the conserved helical domain of the eIF3c-NTD (Fig. 3A, dashed oval) spanning residues 55 through 156. The eIF3c-NTD interacts with the 18S rRNA at the platform region through several evolutionary well-conserved residues on each side of this domain (Ser52, Arg53, Lys56 and Arg127 with A1360, C1361, C1596 and C370, Supplementary Fig. 7O, Table S1), suggesting that it has a similar PIC binding mode also in mammals, despite the obvious differences in binding to eIFs 1 and 5 reported above. In addition to these main contacts with the rRNA, a minor interaction of eIF3c can be observed with eS27 (via residues Glu191 and Lys192 with Glu56 and Lys63) (Fig. 3D, Table S1). In contrast to eIF3c, the eIF3a binding to the ribosomal protein eS1 does not seem to differ from other eukaryotes (residues Thr7, Arg8, Thr12 and Leu17 contact Gln77, Thr72, Arg192 and Ile194, respectively) (Fig. 3C, Table S1).

**Figure 3.**
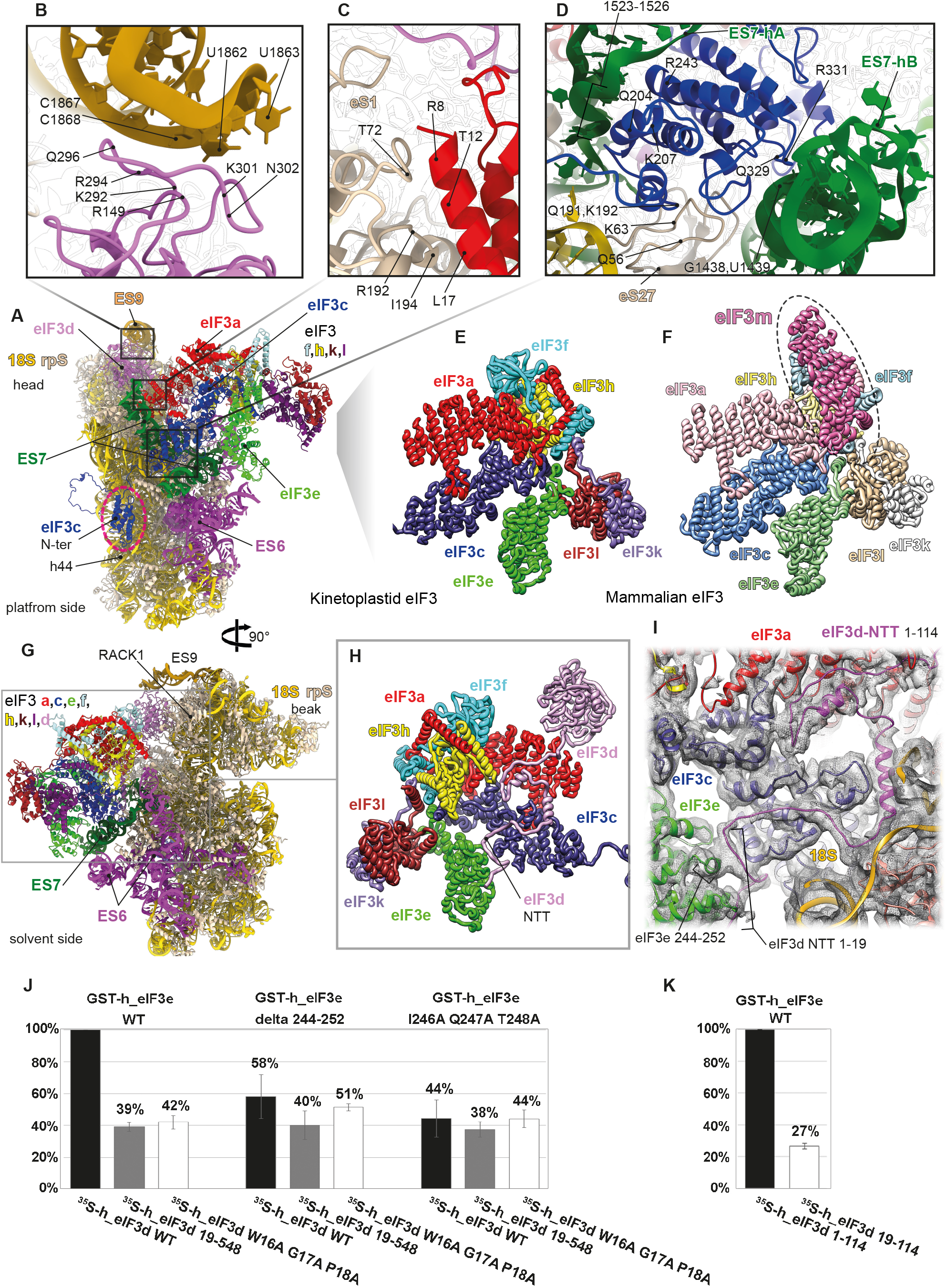
Kinetoplastidian eIF3 and its unique binding site. **(A)** The overall view of the atomic model of the 43S PIC from the platform side. The conserved helical domain of the eIF3c-NTD is encircled with dashed line, eIF3 subunits are colored variably and 18S RNA in yellow. (**B**) Closeup view of the interaction between the ES9S (honey yellow) and eIF3d (in pink). (**C**) Close-up view of the interaction between eIF3a (in red) and eS1 (in beige) (**D**) Close-up view of the interaction between the ES7S (in green) and eIF3c (in blue). (**E**) Cartoon representation of the atomic model of the kinetoplastidian eIF3 structural core. (**F**) Cartoon representation of an atomic model of the mammalian eIF3 structural core. Subunit eIF3m, which is not encoded by kinetoplastids, is marked by dashed oval. (**G**) The overall view of an atomic model of the 43S PIC from the solvent side. (**H**) Cartoon representation of the atomic model of the kinetoplastidian eIF3 focused on the eIF3d-NTT (in pink). (**I**) Fitting of the eIF3d-NTT model into its cryo-EM map. (**J, K**) binding analysis between human eIF3d and GST-eIF3e, expressed in plots showing normalized data from three different dilutions of GST-proteins (see Supplementary Fig. 10A).

Another unusually large ES is the kinetoplastidian ES9^S^ that forms a “horn” on the 40S head, bending towards the mRNA exit channel, where it binds to and stabilizes eIF3d within the 43S PIC (Fig. 3A-B, Table S1), representing another important feature that is specific to translation initiation in trypanosomatids. In particular, the eIF3d main globular domain interacts with ES9^S^ mainly through residues Arg149, Arg294, Gln296, Lys301 and Asp306 contacting nucleotides G1861 through C1867. Moreover, close to the N-terminal tail, eIF3d through Asp43 and Asp50 interacts with G1532 and A1475 (Supplementary Fig. 8 A and B, Table S1). Noteworthy, structures of ES7^S^ and the exceptionally large ES6^S^ (Supplementary Fig. 12) undergo drastic conformational changes upon binding of eIF3, as can be observed by comparing this structure with our previous *T. cruzi* 40S lacking eIF3 (Supplementary Fig. 13B). Amplitude of these conformational acrobatics may indicate their functional importance that, in turn, sets them in the viewfinder for the future drug-targeting studies.

When compared to its mammalian counterpart, the overall conformation of eIF3 structural core differs significantly (Fig. 3E-F, Supplementary 13C-D), mainly due to the lack of the eIF3m subunit in trypanosomatids, which is in part compensated for by the rearrangements of the other core eIF3 subunits like a, c, e, k, l, but mostly f and h. Indeed, eIF3 f and h shift several a-helices and coils to fill for the absence of the m subunit; this rearrangement is probably required for the maintenance of the eIF3 core central helical bundle (Supplementary Fig. 13C-D, arrows indicate the direction of the shift). Moreover, a charge surface analysis reveals very different charge distribution patterns between *T. cruzi* eIF3 and its mammalian counterpart (Supplementary Fig. 14A-B), in part as a consequence of the different 40S binding surface that is mainly represented by rRNA, in contrast to other known eukaryotes.

Importantly, our cryo-EM reconstruction reveals the full structure of eIF3d that appeared separated from the eIF3 structural core in the context of the PIC in all previous studies (Hashem et al., 2013; des Georges et al., 2015; Eliseev et al., 2018). We show here that the eIF3d-NTT, unseen in any previous equivalent complexes, extends towards eIF3e, where it interacts with its PCI domain (residues 1-19 of eIF3d with Ala196, Thr198, Ile 246, Gln247 and Thr248 of eIF3e; Fig. 3G-I, Table S1). Furthermore, the eIF3d-NTT also comes in a less extensive contact with eIF3a, eIF3c and ribosomal protein eS27 (Fig. 3 H and I, Supplementary Fig. 7P-Q and 8C, Table S1). In agreement, the interaction of the eIF3d-NTT (the first 114 residues) with the eIF3 core was previously shown in biochemical and genetics studies (Smith et al., 2016). To support our structural data and investigate the evolutionary conservation of the eIF3d contacts with eIF3 e, a and c subunits within the PIC, we expressed human homologues of all these proteins and subjected them to our GST pull down analysis. As shown in Figures 3 J and K and S10 B and C, the main contact between eIF3d and eIF3e does involve the first 19 residues (in particular W16, G17, and P18) of the former, and residues I246, Q247, and T248 of the latter subunit even in humans. In addition, weak but reproducible binding between eIF3d and eIF3a and eIF3c subunits was also detected, in contrast to other eIF3 subunits (Supplementary Fig. 10D and E). Since human eIF3d was shown to interact with the mRNA cap (Lee et al., 2016) and, together with several other eIF3 subunits (including eIF3a and eIF3e) proposed to promote recruitment of selected mRNAs to the 43S PIC to control their expression in response to various stresses and cellular signals (Lee et al., 2015; Shah et al., 2016), we speculate that these contacts play pivotal role in coordinating the eIF3d-specific functions with the rest of eIF3 on the ribosome.

### The trypanosomatid-specifìc k-DDX60

As mentioned above, our cryo-EM reconstructions of the *T. cruzi* and *L. tarentolae* 43S PICs revealed a large density at the intersubunit side of the 40S (Fig. 1B-D, Supplementary Fig. 6). Known structures of eIFs or ABCE1 (des Georges et al., 2015; Llácer et al., 2018; Erzberger et al., 2014) do not fit into this density and proteomic analysis shows substantial presence of the helicase DDX60 protein in our samples (Fig. 1B, Supplementary Fig. 1 and 2), which we henceforward refer to as kinetoplastidian-DDX60 (k-DDX60). The density was of sufficient resolution to build an atomic model of k-DDX60, including the helicase recombinase A (RecA) domains (Fig. 4, Supplementary Fig. 8 D-H), which fully validates our assignment. Besides the RecA domains, k-DDX60 counts two winged-helices domains, two ratchet domains and one kinetoplastid-specific A-site insert (AI) that protrudes at the end of the RecA2 domain from the C-terminal cassette (Fig. 4C-E, Supplementary Fig. 15 for secondary structures details).

**Figure 4.**
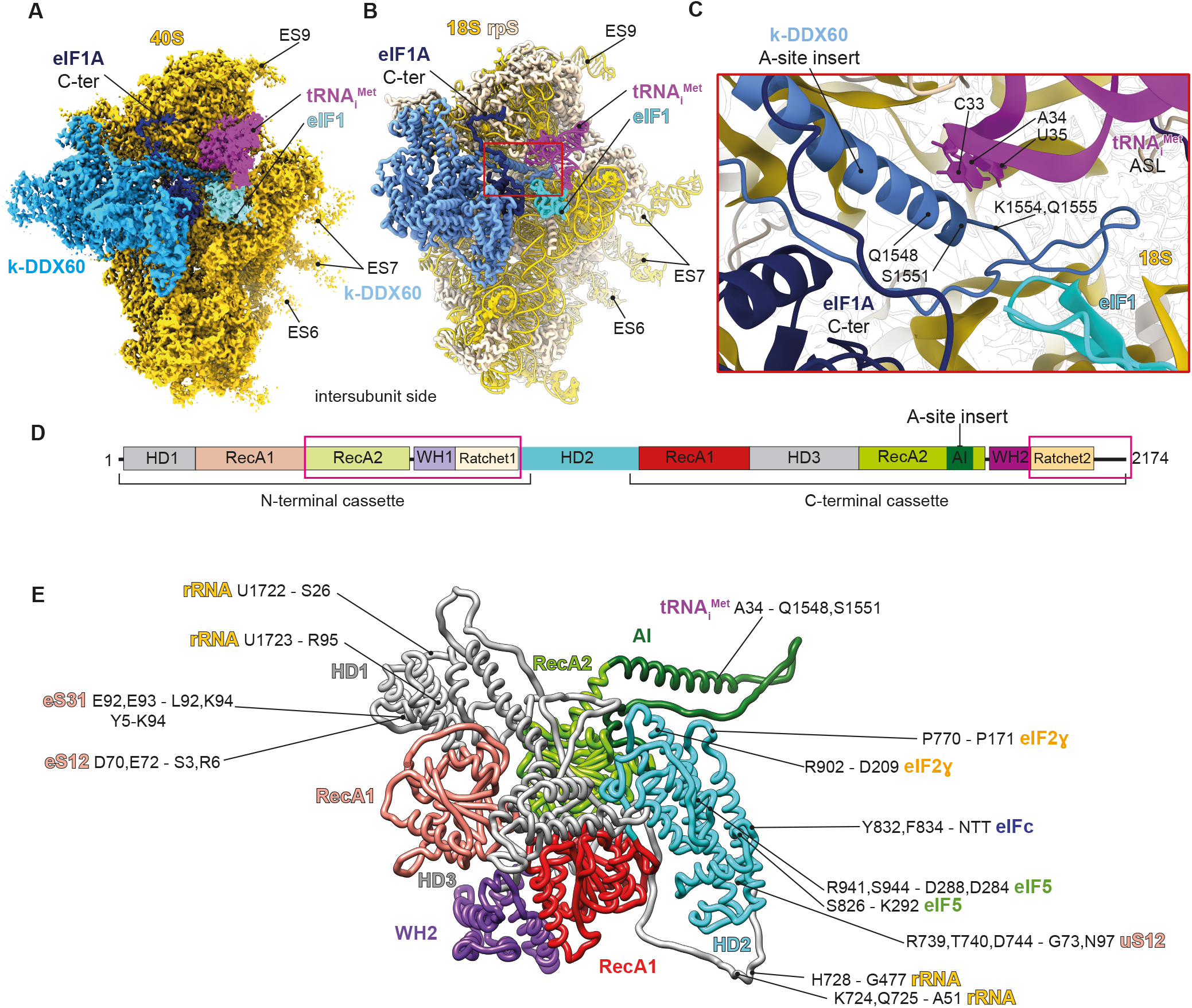
k-DDX60 structure and interactions within the 43S PIC. (**A**) The cryo-EM structure of the *T. cruzi* 43S PIC highlighting k-DDX60 (colored in dark turquoise). eIF 2, 3 and 5 densities were removed for clarity (**B**) Cartoon representation of a partial atomic model of the *T. cruzi* 43S PIC. (**C**) A close-up view of the k-DDX60 A-site insert showing its interaction with the anticodon stem loop (ASL). (**D**) Schematic representation of the k-DDX60 domains. Pink boxes indicate the domains that couldn’t be modeled because of their lower local resolution (See Supplementary Fig. 4). (**E**) Cartoon representation of the atomic model of the k-DDX60 and its interactions with the 43S PIC color-coded in accord with its schematic representation in the panel D.

The presence of k-DDX60 is not due to the use of GMP-PNP, as we did not retrieve any densities resembling GMP-PNP in any of k-DDX60 RecA domains. In addition, its known mammalian DDX60 homologue is an ATP helicase. Next we wanted to inspect structural impact of its ATPase activity by determining the structure of the 43S PIC purified from *T. cruzi* cell lysate supplemented with ATP, in addition to GMP-PNP (Fig. 5A). It is important to stress out that the resolution of the 43S PIC+ATP reconstruction is mostly worse than 4Å, precluding unambiguous determination of whether ATP hydrolysis took place or not. Nonetheless, the structure reveals a global conformational rearrangement of the 40S head (Fig. 5B-C), which could be driven by the k-DDX60 rearrangement upon ATP hydrolysis (Fig. 5D-F). In addition, we also observe the presence of an extra density at the RecA1 domain of the C-terminal cassette at the position that is unoccupied in the absence of ATP (Fig. 5D).

**Figure 5.**
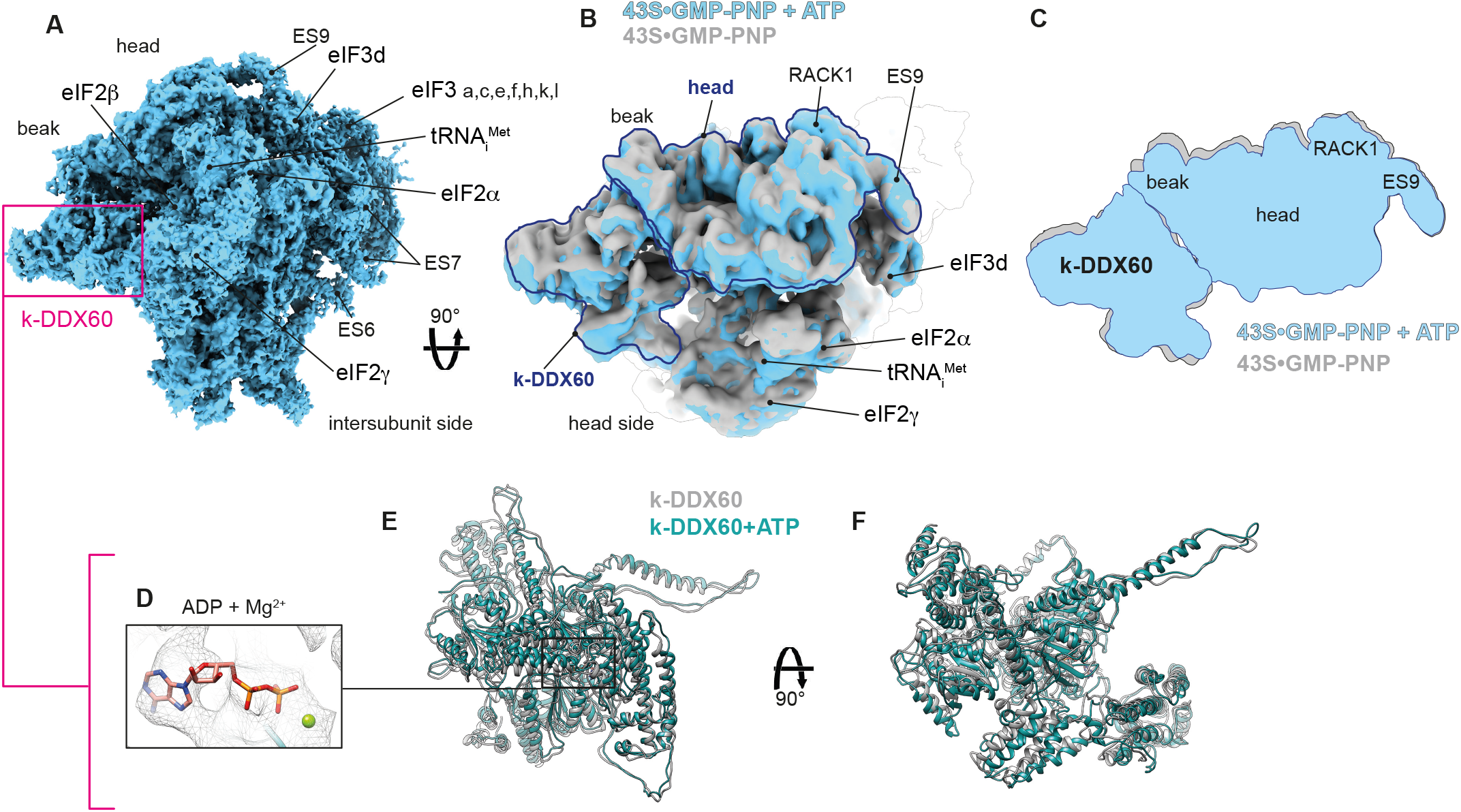
Global conformational rearrangement of the 43S PIC driven by ATP binding to k-DDX60. (**A**) Cryo-EM reconstruction of the *T. cruzi* 43S PIC in the presence of ATP. (**B**) Superposition of the cryo-EM reconstructions of the 43S•GMP-PNP (in grey) and the 43S•GMP-PNP supplemented with ATP (in turquoise), seen from the top. (**C**) Schematic representation of the structural rearrangements induced by ATP. (**D**) A close-up view of the ATP binding pocket within the RecA1 domain of the C-terminal cassette of k-DDX60. (**E, F**) Superimposition of the k-DDX60 atomic model from the cryo-EM structure of the 43S•GMP-PNP and 43S•GMP-PNP supplemented with ATP presented in two different orientations.

k-DDX60 binds both to the head and the body of the 40S and the structural dynamics induced by the ATP addition suggest its involvement in remodeling of the 43S PIC mRNA channel due to the head swiveling. Importantly, the AI extended helix of k-DDX60 interacts with the anticodon stem-loop of the Met-tRNA_i_^Met^ (Fig. 4C, Supplementary Fig. 8P), preventing the codon-anticodon interaction in its presence. The release of k-DDX60, or at least of its AI helix, must therefore precede the rotation of the 40S head and the full accommodation of the Met-tRNA_i_^Met^ in the P-site. Moreover, k-DDX60 interacts directly with eIF3c-NTD and eIF5 (Fig. 4E, Supplementary Fig. 8 I and N), in addition to the 18S rRNA and ribosomal proteins eS12, uS12 and eS31 (Fig. 4E, Supplementary Fig. 8 I-O), suggesting its direct involvement in structural changes accompanying/driving the AUG recognition process. Finally, k-DDX60 comes in close proximity with eIF2β, eIF2g and eIF3c, but the local resolution at these possible interaction sites did not allow to unambiguously define the interacting residues. We believe that owing to its extensive interactions with numerous components of the 43S PIC, k-DDX60 led to a stabilization of the 43S PIC that enabled rigidification of flexible tails of eIFs allowing them to be resolved by cryo-EM. In agreement, most of these interactions occur via additional domains and insertions of k-DDX60 that are inexistent in its mammalian homologue (Fig. 4D, Supplementary Fig. 15B). It is not clear why translation initiation, perhaps in particular the AUG selection process, in kinetoplastids requires this specific helicase. Interestingly, all mature cytoplasmic mRNAs in kinetoplastids possess a 39-nucleotide spliced leader that confers them an unusual hypermethylated 5’-cap structure (known as cap4) (Michaeli, 2011). Therefore, the presence of this helicase might be required for an efficient recruitment and handling of these kinetoplastid-specific mRNAs until the start codon has been recognized.

## CONCLUSIONS

In summary, our structure reveals numerous previously uncharacterized features of the eukaryotic translation initiation machinery, some of which are common to other eukaryotes, such as the placement and proposed roles of terminal tails of eIF1, eIF1A, eIF2β, eIF3c, eIF3d, and, above all, the precise binding site of the eIF5-CTD within the 43S PIC (Fig. 6A-C). Furthermore, our data uncover several striking features of translation initiation specific to kinetoplastids (Fig. 6D-F), such as the role of the oversized kinetoplastidian ES^S^ in providing a large, unique binding surface for eIF3, as well as the structural characterization of k-DDX60. These unique molecular features of translation initiation in kinetoplastids represent an unprecedented opportunity to interfere specifically with the initiation process in these “hard-to-combat” parasites, which may stimulate new venues of research and development of new effective drugs against trypanosomiasis and leishmaniasis.

**Figure 6.**
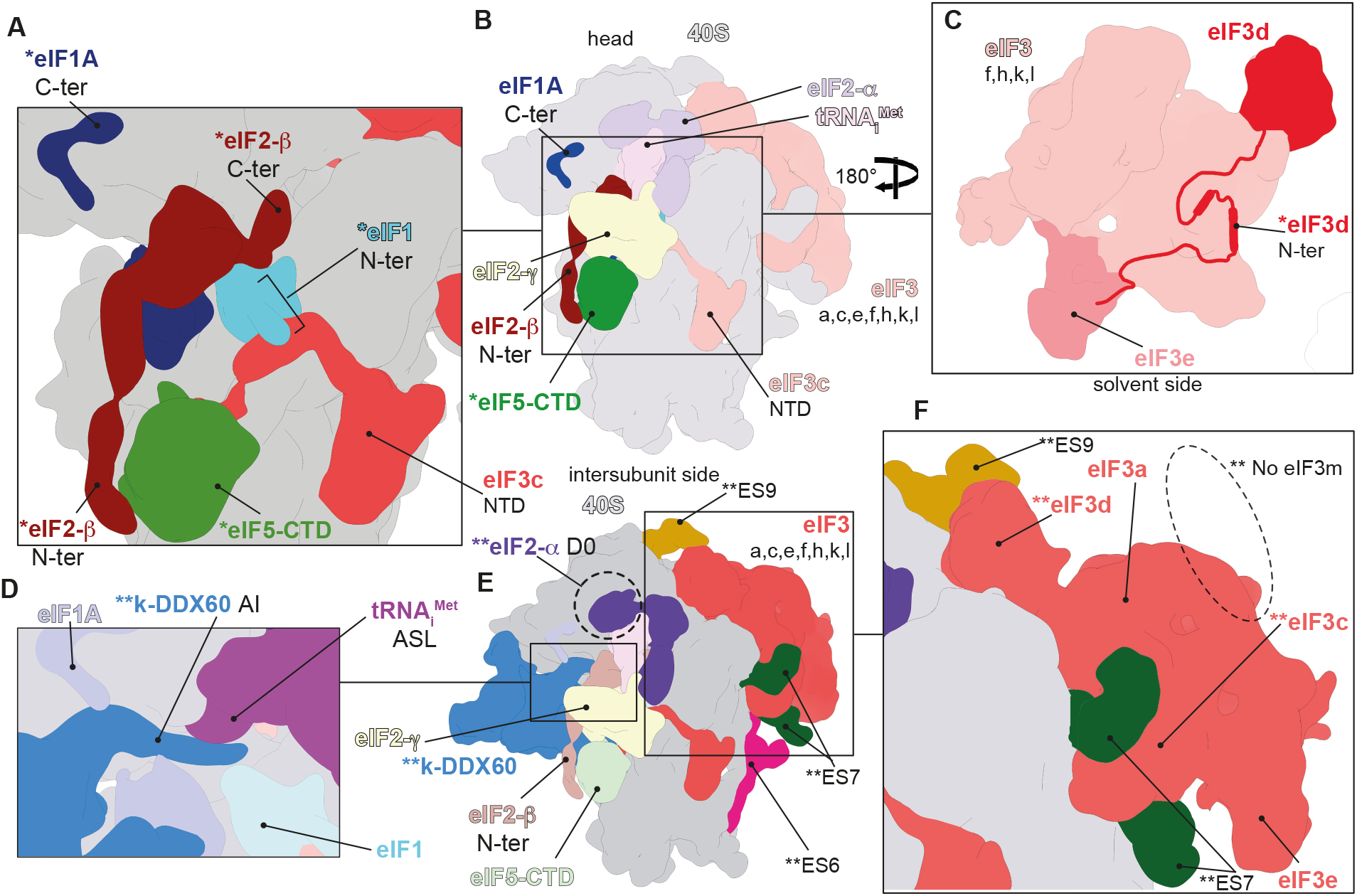
Previously uncharacterized eukaryotic-conserved and trypanosomatid-specific features of the 43S PIC revealed in our work. (**A**) Schematic model representing a close-up view on the N-terminal tails of eIF 1, 1A, 2β, eIF5-CTD and eIF3c-NTD, all conserved among eukaryotes and revealed in the current work. The ternary complex was removed for clarity. (**B**) Schematic model representing the 43S PIC from the intersubunit side. The previously uncharacterized features revealed in our work are colored in brighter colors. (**C**) Schematic model representing a solvent side view of eIF3 highlighting the conserved N-terminal tail of eIF3d and its main interactions with eIF3e, revealed in the current work. (**D**) Schematic model representing a close-up view on the A-site Insert of k-DDX60 and its interaction with the anti-codon stem-loop (ASL). (**E**) Schematic model representing the *T. cruzi* 43S PIC from the intersubunit side. Dashed circle highlights the kinetoplastid-specific domain eIF2α, dubbed here “D0”. The kinetoplastid-specific features revealed in our work are colored in brighter colors. (**F**) Schematic model representing a close-up view on the kinetoplastidian eIF3 showing its specific interaction with ES7^S^ and ES9^S^, and the absence of the eIF3m subunit. *=Conserved features among eukaryotes revealed in our work. **=Kinetoplastid-specific features revealed in our work.

## ACKNOWLEDGMENTS

We thank Christoph Diebolder, and Ludovic Renault (NeCEN, Leiden, Holland) as well as Julio Ortiz Espinoza, Corinne Crucifix and and Christine Ruhlmann (IGBMC, Strasbourg, France) for assistance with data acquisition. We would like also to thank Eder Mancera-Martinez for her help in the sample’s purification and the High-Performance Computing Center of the University of Strasbourg for IT support. The mass spectrometry instrumentation was funded by the University of Strasbourg, IdEx “Equipement mi-lourd” 2015.

## AUTHOR CONTRIBUTIONS

J.B.Q. and A.S. purified the complexes from *T. cruzi*, M.L.D.C. and A.R.R. purified the complex from *L. tarentolae*. T.P. preformed the GST pulldown assays and analyzed the data together with L.S.V. Y.H. and H.S. performed the cryo-EM data processing. L.K. performed MS/MS analysis. A.B., J.B.Q. and Y.H. interpreted the cryo-EM data. A.B. and Y.H. performed the molecular modeling. J.B.Q., T.P., A.B., L.S.V. and Y.H. wrote the manuscript. Y.H. supervised the research.

## AUTHOR INFORMATION

The cryo-EM maps of the 43S+GMPPNP, 43S+GMPPNP+ATP PICs from *T. cruzi* and the 43S+GMPPNP PIC from *L. tarentolae* have been deposited at the Electron Microscopy Data Bank (EMDB) with accession codes EMD-11893, EMD-11895 and EMD-11896. The atomic model of the 43S+GMPPNP PIC from *T. cruzi* has been deposited in the Protein Data Bank (PDB) with the accession code 7ASE. K-DDX60 atomic model was fitted in its density from the 43S+GMPPNP+ATP PIC, with the accession code 7ASK.

The authors declare no competing financial interests. Correspondence and requests for materials should be addressed to Y.H.

## METHODS DETAILS

### Kinetoplastids Cultures

*Trypanosoma cruzi* epimastigoes (Y strain - TcII) were grown at 28°C in liver infusion tryptose (LIT) medium, supplemented with 10% heat-inactivated fetal bovine serum. *Leishmania tarentolae* strain T7-TR (Jena Bioscience) were grown at 26°C in brain-heart infusion-based medium (LEXSY BHI; Jena Bioscience), supplemented with Nourseothricin and LEXSY Hygro (Jena Bioscience), hemin and penicillin-streptomycin.

### 48S Initiation Complex Purification

*T. cruzi* and *L. tarentolae* 48S initiation complexes were grown to a density 3·10^6^ per mL and 2.5·10^6^ per mL, for *T. cruzi* and *L. tarantolae*, respectively, in 200 mL flasks in culture medium. The parasites were harvested, put in buffer I (20 mM HEPES-KOH pH 7.4, 100 mM KOAc, 4 mM Mg (OAc)2, 2 mM DTT, EDTA free protease inhibitor cocktail and RNasin inhibitor) and subjected to lysis by freeze-thaw cycles. After the centrifugation at 12,000 *g* for 30 min at 4°C, the supernatant was incubated in the presence of 10 mM GMP-PNP (the non-hydrolyzable analog of GTP) for 10 min at 28°C. The supernatant was layered onto 10-30 % (w/v) sucrose gradients and centrifuged (35 000 rpm, 5h30 min, 4°C) using an SW41 Ti rotor (Beckman-Coulter). The fractions containing 48S ICs were collected and pooled according the UV absorbance profile. Buffer was exchanged by precipitating ribosomal complexes and re-suspending them in sucrose-free buffer II (10 mM HEPES-KOH pH 7.4, 50 mM KOAc, 10 mM NH4Cl, 5 mM Mg(OAc)2, and 2 mM DTT). For the ATP supplemented 43S PIC, the protocol above was repeated for *T. cruzi* with an addition of 10 mM of ATP.

### Cryo-EM Grid preparation

Grid preparation: 4 μL of the sample at a concentration of 90 nM was applied onto the Quantifoil R2/2 300-mesh holey carbon grid, which had been coated with thin carbon film (about 2nm) and glow-discharged. The sample was incubated on the grid for 30 sec and then blotted with filter paper for 1.5 sec in a temperature and humidity controlled Vitrobot Mark IV (T = 4°C, humidity 100%, blot force 5) followed by vitrification in liquid ethane.

### Cryo-EM Image acquisition

Data collections of the three described molecular complexes were performed on three different instruments. The main complex (*T. cruzi* 43S PIC) was imaged (at the IGBMC EM facility, Illkirch, France) on a spherical aberration corrected Titan Krios S-FEG instrument (FEI Company) at 300 kV using the EPU software (Thermo Fisher Company) for automated data acquisition. Data were collected at a nominal under focus of −0.6 to −4.5 μm at a magnification of 127,272 X yielding a pixel size of 1.1 Å. Micrographs were recorded as movie stack on a Gatan Summit K2 direct electron detector, each movie stack were fractionated into 20 frames for a total exposure of an electron dose of 30 ē/Å^2^. The *T. cruzi* 43S PIC supplemented with ATP was imaged with the exact setup described above, but in the Netherland’s NeCEN EM facility, Leiden, which is not Cs corrected. The *L. tarentolae* 43S PIC dataset was collected (at the IECB EM facility, Pessac, France) on a Talos Artica instrument (FEI Company) at 200 kV using the EPU software (FEI Company) for automated data acquisition. Data were collected at a nominal underfocus of −0.5 to −2.7 μm at a magnification of 120,000 X yielding a pixel size of 1.21 Å. Micrographs were recorded as movie stack on a Falcon III direct electron detector (FEI Compagny), each movie stack were fractionated into 20 frames for a total exposure of 1 sec corresponding to an electron dose of 40 ē/Å2.

### Image processing

For all three datasets, drift and gain correction and dose weighting were performed using MotionCor2 (Zheng et al., 2017). A dose weighted average image of the whole stack was used to determine the contrast transfer function with the software Gctf (Zhang et al., 2016). The following process has been achieved using RELION 3.0 (Zivanov et al., 2018). Particles were picked using a Laplacian of gaussian function (min diameter 300 Å, max diameter 320 Å). For the main dataset (*T. cruzi* 43S PIC), particles were then extracted with a box size of 360 pixels and binned threefold for 2D classification into 200 classes, yielding 202,920 particles presenting 40S-like shape. These particles were then subjected to 3D classification into 10 classes. Two subclasses depicting high-resolution and 48S features have been selected for a second round of classification into two classes. One class ended as a possible 48S complex (12910 particles, don’t present densities for k-DDX60) and a second as a 43S+DDX60 complex (33775 particles). Refinement of the 43S-DDX60 complex yielded an average resolution of 3.3Å. The 48S class was not analyzed any further. Determination of the local resolution of the final density map was performed using ResMap (Kucukelbir et al., 2014). The dataset of the *T. cruzi* 43S PIC supplemented with ATP was processed identically. However, the sample was more diluted compared to the above-described main complex, thus yielding less particles count after the first 2D classification (98,840 particles presenting 40-like shape). Following the similar classification/processing fashion and after 3D classification, 19700 particles were used to reconstruction a ~4.3Å 43S PIC bound to ATP. Finally, the *L. tarentolae* 43S PIC dataset was processed also identically to the protocol described above. As the aim of this reconstruction is simply to validate the conservation of the architecture in leishmania, only a small dataset was collected, which after processing only yielded ~10,000 particles that were then used to reconstruct the 43S PIC at 8.1Å.

### Figure preparation

Figures featuring cryo-EM densities as well as atomic models were visualized with UCSF Chimera (Pettersen et al., 2004).

### Mass spectrometry analysis and data post-processing

Protein extracts were precipitated overnight with 5 volumes of cold 0.1 M ammonium acetate in 100% methanol. Proteins were then digested with sequencing-grade trypsin (Promega, Fitchburg, MA, USA) as described previously^*5*^. Each sample was further analyzed by nanoLC-MS/MS on a QExactive+ mass spectrometer coupled to an EASY-nanoLC-1000 (Thermo-Fisher Scientific, USA). Peptides and proteins were identified with Mascot algorithm (version 2.5.1, Matrix Science, London, UK) and data were further imported into Proline v1.4 software (http://proline.profiproteomics.fr/). Proteins were validated on Mascot pretty rank equal to 1, and 1% FDR on both peptide spectrum matches (PSM score) and protein sets (Protein Set score). The total number of MS/MS fragmentation spectra was used to relatively quantify each protein (Spectral Count relative quantification). Proline was further used to align the Spectral Count values across all samples. The whole MS dataset was then normalized.

The mass spectrometric data were deposited to the ProteomeXchange Consortium via the PRIDE partner repository with the dataset identifier PXD016063 (Reviewer account details, rhv9KZXk).

### Volcano plot

Volcano plot presented in Fig. 1 was obtained after manual validation of the results. For that end, we only consider proteins that present at least 5 spectra. Further validation was performed by analysing the pre-initiation complex after further purification step using size exclusion chromatography.

### Model building and refinement

The atomic model of the preinitiation complex 48S from Trypanosoma *cruzi* was built using the modelling softwares Chimera (Pettersen et al., 2004), Coot (Emsley et al., 2004), Phenix (Adams et al., 2010) and VMD (Humphrey et al., 1996).

The previous 40S structure of Trypanosoma *cruzi* (Brito Querido et al., 2017) (PDBID : 5OPT) was used to build the core of the initiation complex containing the small subunit ribosomal RNA and proteins. The head required a rotation to fit the new structure.

The ternary complex (tRNA, eIF2α, eIF2γ), eIF2β, eIF1a and eIF1 were thread from the translation initiation complex of yeast (Llácer et al., 2015) (PDBID : 3JAQ).

DDX60-like starting point was the recA domains from the human helicase protein Brr2 (Santos et al., 2012) (PDBID : 4F93). The remaining domains of DDX60-like was built *ab initio* using Coot modelling tools and Chimera “build structure” tools with the help of sympred (Simossis et al., 2004) for secondary structure prediction and the homology modelling webservices Swissmodel (Waterhouse et al., 2018) and phyre2 (Kelley et al., 2015).

eIF3 was thread from the already published mammalian eIF3 (des Georges et al., 2015) (PDBID : 5A5T), subunit m was deleted since it’s not present in Kinetoplastid and rearrangements of the nearby subunits were made. Subunit d was thread from the eIF3d crystal structure of Nasonia vitripennis (Lee et al., 2016) (PDBID : 5K4B) and the N-terminal tail was built in Chimera.

eIF5 Cter-domain was thread from the eIF5 crystal from human (Bieniossek et al., 2006) (PDBID : 2IU1).

The global atomic model was refined using the Molecular Dynamic Flexible Fitting (Trabuco et al., 2008) then the geometry parameters were corrected using PHENIX real space refine for proteins and eraser (Chou et al., 2012) for RNA.

### Secondary structures of k-DDX60 and the 18S

The secondary structure of the 18S was done based on the *S.c*. 18S template downloaded from the RiboVision Webservice (Bernier et al., 2014). The secondary structures of the 18S expansion segments were edited manually based on the 3D atomic model of the complex. The sequence and residues numbering were corrected consistently with *T. cruzi*.

The secondary structure of k-DDX60 was derived from its 3D atomic model (this work) using the PDBsum Webservice (Laskowski et al., 2018)

### GST pulldown assay

Glutathione S-transferase (GST) pull down experiments with GST fusions and *in vitro* synthesized ^35^S-labeled polypeptides were conducted as described previously (PMID:11179233). Briefly, individual GST-fusion proteins were expressed in *Escherichia coli* (BL-21 Star DE3 or BL21 Rosett2 DE3). Bacterial culture was grown at 37°C in the LB medium to OD 0.6-0.8 and the synthesis of GST-fusion proteins were induced by the addition of 1mM IPTG. After 2 hr of shaking at 37°C or overnight at 16°C the cells were harvested, resuspended in a Phosphate-buffered saline (PBS), and subjected to mechanical lysis with a subsequent agitation in the presence of 1-1.5% Triton X-100 for 30 min at 4°C. The GST-proteins were then immobilized on glutathione sepharose beads (GE Healthcare, cat # GE17-0756-01) from the pre-cleaned supernatant, followed by three washing steps with the 1 ml of phosphate buffered saline.^35^S-labeled polypeptides were produced *in-vitro* by the TnT^®^ Quick Coupled Transcription/Translation System (Promega cat # L1170) according to the vendor’s instructions.

To examine the binding, individual GST fusions were incubated with ^35^S-labeled proteins at 4°C for 2 h in buffer B (20mM HEPES (pH 7,5), 75mM KCl, 0,1mM EDTA, 2,5mM MgCl2, 0,05% IGEPAL, 1mM DTT). For experiments requiring more stringent conditions the buffer B was supplement with 1% fat free milk. Subsequently, the beads were washed three times with 1 ml of phosphate buffered saline and interacting proteins were separated by SDS-PAGE. Gels were first stained with Gelcode Blue stain reagent (Thermofisher, cat # 24592) and then subjected to autoradiography.

Quantification of binding experiments was done by the Quantity One software. The data was generated as an adjusted volume with the local background subtraction and linear regression methods. The data for each ^35^S-labeled protein was first normalized to its input and the percentage of input binding was then calculated. The resulting data was subsequently normalized to its corresponding control (for Fig. 3J: ^35^S-eIF3d WT – GST-eIF3e WT; and for Fig. 3K: ^35^S-eIF3d 1-114 – GST-eIF3e WT) and means from three different dilutions of GST-fusions were calculated; errors bars indicate standard deviation.

## Plasmid constructions

See supplementary text.

## REFERENCES

P. D. Adams, P. V. Afonine, G. Bunkóczi, V. B. Chen, I. W. Davis, N. Echols, et al., PHENIX: a comprehensive Python-based system for macromolecular structure solution. Acta Cryst. D66, 213–221 (2010).

P. V. Alone, T. E. Dever, Direct binding of translation initiation factor eIF2gamma-G domain to its GTPase-activating and GDP-GTP exchange factors eIF5 and eIF2B epsilon. J. Biol. Chem. 281, 12636–44 (2006).

M. A. Algire, D. Maag, J. R. Lorsch, Pi release from eIF2, not GTP hydrolysis, is the step controlled by start-site selection during eukaryotic translation initiation. Mol. Cell. 20, 251–62 (2005).

S. Alsford, D. J. Turner, S. O. Obado, A. Sanchez-Flores, L. Glover, M. Berriman, C. Hertz-Fowler, D. Horn, High-throughput phenotyping using parallel sequencing of RNA interference targets in the African trypanosome. Genome Res. 21, 915–24 (2011).

K. Asano, T. Krishnamoorthy, L. Phan, G. D. Pavitt, A. G. Hinnebusch, Conserved bipartite motifs in yeast eIF5 and eIF2Bepsilon, GTPase-activating and GDP-GTP exchange factors in translation initiation, mediate binding to their common substrate eIF2. EMBO J. 18, 1673–88 (1999).

K. Asano, A. Shalev, L. Phan, K. Nielsen, J. Clayton, L. Valásek, T. F. Donahue, A. G. Hinnebusch, Multiple roles for the C-terminal domain of eIF5 in translation initiation complex assembly and GTPase activation. EMBO J. 20, 2326–37 (2001).

C. R. Bernier, A. S. Petrov, C. C. Waterbury, J. Jett, F. Li, L. E. Freil, X. Xiong, L. Wang, B. L. R. Migliozzi, E. Hershkovits, Y. Xue, C. Hsiao, J. C. Bowman, S. C. Harvey, M. A. Grover, Z. J. Wartell, and L. D. Williams. FD169: RiboVision Suite for Visualization and Analysis of Ribosomes. Faraday Discussions (2014).

C. Bieniossek, P. Schutz, M. Bumann, A. Limacher, I. Uson, U. Baumann, The Crystal Structure of the Carboxy-Terminal Domain of Human Translation Initiation Factor Eif5. J.Mol.Biol. 360, 457 (2006).

J. Brito Querido, E. Mancera-Martínez, Q. Vicens, A. Bochler, J. Chicher, A. Simonetti, Y. Hashem, The cryo-EM Structure of a Novel 40S Kinetoplastid-Specific Ribosomal Protein. Structure. 25, 1785–1794.e3 (2017).

F. C. Chou, P. Sripakdeevong, S. M. Dibrov, T. Hermann, and R. Das, Correcting pervasive errors in RNA crystallography through enumerative structure prediction. Nat Methods 10, 74–76 (2012).

S. Das, T. Maiti, K. Das, U. Maitra, Specific interaction of eukaryotic translation initiation factor 5 (eIF5) with the beta-subunit of eIF2. J. Biol. Chem. 272, 31712–31718 (1997).

S. Das, and U. Maitra. Mutational analysis of mammalian translation initiation factor 5 (eIF5): role of interaction between the beta subunit of eIF2 and eIF5 in eIF5 function in vitro and in vivo. Mol. Cell. Biol. 20, 3942–3950 (2000).

A. des Georges, V. Dhote, L. Kuhn, C. U. T. Hellen, T. V Pestova, J. Frank, Y. Hashem, Structure of mammalian eIF3 in the context of the 43S preinitiation complex. Nature. 525, 491–5 (2015).

B. Eliseev, L. Yeramala, A. Leitner, M. Karuppasamy, E. Raimondeau, K. Huard, E. Alkalaeva, R. Aebersold, C. Schaffitzel, Structure of a human cap-dependent 48S translation pre-initiation complex. Nucleic Acids Res. 46, 2678–2689 (2018).

P. Emsley and K. Cowtan, Coot: model-building tools for molecular graphics. Acta Crystallogr. D60, 2126–2132 (2004).

J. P. Erzberger, F. Stengel, R. Pellarin, S. Zhang, T. Schaefer, C. H. S. Aylett, P. Cimermančič, D. Boehringer, A. Sali, R. Aebersold, N. Ban, Molecular Architecture of the 40S ▪ eIF1 ▪ eIF3 Translation Initiation Complex. Cell. 158, 1123–1135 (2014).

C. S. Fraser, J. Y. Lee, G. L. Mayeur, M. Bushell, J. A. Doudna, J. W. Hershey. The j-subunit of human translation initiation factor eIF3 is required for the stable binding of eIF3 and its subcomplexes to 40S ribosomal subunits in vitro. J Biol Chem. 279, 8946–8956 (2004).

E. Guca and Y. Hashem, Major structural rearrangements of the canonical eukaryotic translation initiation complex. Curr. Opin. Struct. Biol. 53, 151–158 (2018).

A. Simonetti, E. Guca, A. Bochler, L. Kuhn, Y. Hashem. Structural insights into the mammalian late-stage initiation complexes. Cell Rep. 31, 107497 (2020).

Y. Hashem, A. des Georges, V. Dhote, R. Langlois, H. Y. Liao, R. A. Grassucci, C. U. T. Hellen, T. V Pestova, J. Frank, Structure of the mammalian ribosomal 43S preinitiation complex bound to the scanning factor DHX29. Cell. 153, 1108–19 (2013).

Y. Hashem, A. des Georges, J. Fu, S. N. Buss, F. Jossinet, A. Jobe, Q. Zhang, H. Y. Liao, R. A. Grassucci, C. Bajaj, E. Westhof, S. Madison-Antenucci, J. Frank, High-resolution cryoelectron microscopy structure of the Trypanosoma brucei ribosome. Nature. 494, 385–9 (2013).

Y. Hashem, J. Frank, The Jigsaw Puzzle of mRNA Translation Initiation in Eukaryotes: A Decade of Structures Unraveling the Mechanics of the Process. Annu. Rev. Biophys. (2018), doi:10.1146/annurev-biophys-070816-034034.

A. Hinnebusch, Structural Insights into the Mechanism of Scanning and Start Codon Recognition in Eukaryotic Translation Initiation. Trends Biochem. Sci. 42, 589–611 (2017).

W. Humphrey, A. Dalke and K. Schulten, VMD - Visual Molecular Dynamics. J. Molec. Graphics 14, 33–38 (1996).

M. Karásková, S. Gunišová, A. Herrmannová, S. Wagner, V. Munzarová, L. S. L. S. Valášek, S. Gunisová, L. S. Valásek, Functional Characterization of the Role of the N-terminal Domain of the c/Nip1 Subunit of Eukaryotic Initiation Factor 3 (eIF3) in AUG Recognition. J. Biol. Chem. 287, 28420–34 (2012).

K. Kashiwagi, T. Yokoyama, M. Nishimoto, M. Takahashi, A. Sakamoto, M. Yonemochi, M. Shirouzu, T. Ito, Structural basis for eIF2B inhibition in integrated stress response. Science. 364, 495–499 (2019).

L. A. Kelley, S. Mezulis, C. M. Yates, M. N. Wass, M. J. E. Sternberg, The Phyre2 web portal for protein modeling, prediction and analysis. Nature Protocols 10, 845–858 (2015)

L. R. Kenner, A. A. Anand, H. C. Nguyen, A. G. Myasnikov, C. J. Klose, L. A. McGeever, J. C. Tsai, L. E. Miller-Vedam, P. Walter, A. Frost, eIF2B-catalyzed nucleotide exchange and phosphoregulation by the integrated stress response. Science. 364, 491–495 (2019).

A. Kucukelbir, F. J. Sigworth and H. D. Tagare. “Quantifying the local resolution of cryo-EM density maps.” Nat Methods 11, 63–6551 (2014).

R. A. Laskowski, J. Jabłońska, L. Pravda, R. S. Vařeková, J. M. Thornton. PDBsum: Structural summaries of PDB entries. Prot. Sci. 27, 129–134 (2018).

J. L. Llácer, T. Hussain, L. Marler, C. E. Aitken, A. Thakur, J. R. Lorsch, A. G. Hinnebusch, V. Ramakrishnan, Conformational Differences between Open and Closed States of the Eukaryotic Translation Initiation Complex. Mol. Cell. 59, 399–412 (2015).

J. L. Llácer, T. Hussain, A. K. Saini, J. S. Nanda, S. Kaur, Y. Gordiyenko, R. Kumar, A. G. Hinnebusch, J. R. Lorsch, V. Ramakrishnan, Translational initiation factor eIF5 replaces eIF1 on the 40S ribosomal subunit to promote start-codon recognition. Elife. 7 (2018), doi:10.7554/eLife.39273.

A. S. Y. Lee, P. J. Kranzusch, J. H. D. Cate, eIF3 targets cell-proliferation messenger RNAs for translational activation or repression. Nature (2015), doi:10.1038/nature14267.

A. S. Lee, P. J. Kranzusch, J. A. Doudna, J. H. D. Cate, eIF3d is an mRNA cap-binding protein that is required for specialized translation initiation. Nature. 536, 96–9 (2016).

K. Li, S. Zhou, Q. Guo, X. Chen, D. Lai, Z. Lun, X. Guo, The eIF3 complex of Trypanosoma brucei: composition conservation does not imply the conservation of structural assembly and subunits function. RNA. 23, 333–345 (2017).

R. E. Luna, H. Arthanari, H. Hiraishi, J. Nanda, P. Martin-Marcos, M. A. Markus, B. Akabayov, A. G. Milbradt, L. E. Luna, H.-C. Seo, S. G. Hyberts, A. Fahmy, M. Reibarkh, D. Miles, P. R. Hagner, E. M. O’Day, T. Yi, A. Marintchev, A. G. Hinnebusch, J. R. Lorsch, K. Asano, G. Wagner, The C-terminal domain of eukaryotic initiation factor 5 promotes start codon recognition by its dynamic interplay with eIF1 and eIF2β. Cell Rep. 1, 689–702 (2012).

S. Michaeli. Trans-splicing in trypanosomes: machinery and its impact on the parasite transcriptome. Future Microbiol. 6, 459–474 (2011).

M. Miyashita, H. Oshiumi, M. Matsumoto, T. Seya, DDX60, a DEXD/H box helicase, is a novel antiviral factor promoting RIG-I-like receptor-mediated signaling. Mol. Cell. Biol. 31, 3802–19 (2011).

E. Obayashi, R. E. Luna, T. Nagata, P. Martin-Marcos, H. Hiraishi, C. R. Singh, J. P. Erzberger, F. Zhang, H. Arthanari, J. Morris, R. Pellarin, C. Moore, I. Harmon, E. Papadopoulos, H. Yoshida, M. L. Nasr, S. Unzai, B. Thompson, E. Aube, S. Hustak, F. Stengel, E. Dagraca, A. Ananbandam, P. Gao, T. Urano, A. G. Hinnebusch, G. Wagner, K. Asano, Molecular Landscape of the Ribosome Pre-initiation Complex during mRNA Scanning: Structural Role for eIF3c and Its Control by eIF5. Cell Rep. 18, 2651–2663 (2017).

H. Oshiumi, M. Miyashita, M. Okamoto, Y. Morioka, M. Okabe, M. Matsumoto, T. Seya, DDX60 Is Involved in RIG-I-Dependent and Independent Antiviral Responses, and Its Function Is Attenuated by Virus-Induced EGFR Activation. Cell Rep. 11, 1193–207 (2015).

E. F. Pettersen, T. D. Goddard, C. C. Huang, G. S. Couch, D. M. Greenblatt, E. C. Meng, T. E. Ferrin, UCSF Chimera--a visualization system for exploratory research and analysis. J Comput Chem 25, 1605–12 (2004).

L. Phan, X. Zhang, K. Asano, J. Anderson, H. P. Vornlocher, J. R. Greenberg, J. Qin, A. G. Hinnebusch, Identification of a translation initiation factor 3 (eIF3) core complex, conserved in yeast and mammals, that interacts with eIF5. Mol. Cell. Biol. 18, 4935–46 (1998).

A. M. Rezende, L. A. Assis, E. C. Nunes, T. D. da Costa Lima, F. K. Marchini, E. R. Freire, C. R. S. Reis, O. P. de Melo Neto, The translation initiation complex eIF3 in trypanosomatids and other pathogenic excavates--identification of conserved and divergent features based on orthologue analysis. BMC Genomics. 15, 1175 (2014).

K. F. Santos, S. M. Jovin, G. Weber, V. Pena, R. Luhrmann, M. C. Wahl, Structural basis for functional cooperation between tandem helicase cassettes in Brr2-mediated remodeling of the spliceosome. Proc.Natl.Acad.Sci.USA 109: 17418–17423 (2012).

A. Simonetti, J. Brito Querido, A. G. Myasnikov, E. Mancera-Martinez, A. Renaud, L. Kuhn, Y. Hashem, eIF3 Peripheral Subunits Rearrangement after mRNA Binding and Start-Codon Recognition. Mol. Cell. 63, 206–217 (2016).

V. A. Simossis and J. Heringa, Optimally segmented consensus secondary structure prediction. Bioinformatics (2004).

M. Shah, D. Su, J. S. Scheliga, T. Pluskal, S. Boronat, K. Motamedchaboki, A. R. Campos, F. Qi, E. Hidalgo, M. Yanagida, D. A. Wolf, A Transcript-Specific eIF3 Complex Mediates Global Translational Control of Energy Metabolism. Cell Rep. 16, 1891–902 (2016).

C. R. Singh, Y. Yamamoto, K. Asano, Physical association of eukaryotic initiation factor (eIF) 5 carboxyl-terminal domain with the lysine-rich eIF2beta segment strongly enhances its binding to eIF3. J. Biol. Chem. 279, 49644–55 (2004).

C. R. Singh, R. Watanabe, W. Chowdhury, H. Hiraishi, M. J. Murai, Y. Yamamoto, D. Miles, Y. Ikeda, M. Asano, K. Asano, Sequential eukaryotic translation initiation factor 5 (eIF5) binding to the charged disordered segments of eIF4G and eIF2β stabilizes the 48S preinitiation complex and promotes its shift to the initiation mode. Mol. Cell. Biol. 32, 3978–89 (2012).

D. B. Smith, K. S. Johnson, Single-step purification of polypeptides expressed in Escherichia coli as fusions with glutathione S-transferase. Gene. 67, 31–40 (1988).

M. D. Smith, L. Arake-Tacca, A. Nitido, E. Montabana, A. Park, J. H. Cate, Assembly of eIF3 Mediated by Mutually Dependent Subunit Insertion. Structure. 24, 886–96 (2016).

E. Stolboushkina, S. Nikonov, A. Nikulin, U. Bläsi, D. J. Manstein, R. Fedorov, M. Garber, O. Nikonov, Crystal structure of the intact archaeal translation initiation factor 2 demonstrates very high conformational flexibility in the alpha- and beta-subunits. J. Mol. Biol. 382, 680–91 (2008).

C. Sun, A. Todorovic, J. Querol-Audí, Y. Bai, N. Villa, M. Snyder, J. Ashchyan, C. S. Lewis, A. Hartland, S. Gradia, C. S. Fraser, J. A. Doudna, E. Nogales, J. H. D. Cate, Functional reconstitution of human eukaryotic translation initiation factor 3 (eIF3). Proc. Natl. Acad. Sci. U. S. A. 108, 20473–8 (2011).

A. Thakur, L. Marler, A. G. Hinnebusch, A network of eIF2β interactions with eIF1 and Met-tRNAi promotes accurate start codon selection by the translation preinitiation complex. Nucleic Acids Res. 47, 2574–2593 (2019).

L. G. Trabuco, E. Villa, K. Mitra, J. Frank, and K. Schulten, Flexible fitting of atomic structures into electron microscopy maps using molecular dynamics. Structure 16, 673–683 (2008).

L. Valášek, H. Trachsel, J. Hašek, H. Ruis, Rpg1, the Saccharomyces cerevisiae homologue of the largest subunit of mammlian translation initiation factor 3, is required for translational activity. J. Biol. Chem. 273, 21253–21260 (1998).

L. Valásek, K. H. Nielsen, F. Zhang, C. A. Fekete, A. G. Hinnebusch, Interactions of eukaryotic translation initiation factor 3 (eIF3) subunit NIP1/c with eIF1 and eIF5 promote preinitiation complex assembly and regulate start codon selection. Mol. Cell. Biol. 24, 943–755 (2004).

L. S. Valásek, ‘Ribozoomin‘--translation initiation from the perspective of the ribosome-bound eukaryotic initiation factors (eIFs). Curr. Protein Pept. Sci. 13, 305–30 (2012).E.

L. S. Valášek, J. Zeman, S. Wagner, P. Beznosková, Z. Pavlíková, M. P. Mohammad, V. Hronová, A. Herrmannová, Y. Hashem, S. Gunišová, Embraced by eIF3: structural and functional insights into the roles of eIF3 across the translation cycle. Nucleic Acids Res. 45, 10948–10968 (2017).

S. Wagner, A. Herrmannová, R. Malík, L. Peclinovská, L. S. Valášek, Functional and biochemical characterization of human eukaryotic translation initiation factor 3 in living cells. Mol. Cell. Biol. 34, 3041–52 (2014).

S. Wagner, A. Herrmannová, D. Šikrová, L. S. Valášek, Human eIF3b and eIF3a serve as the nucleation core for the assembly of eIF3 into two interconnected modules: the yeast-like core and the octamer. Nucleic Acids Res. (2016), doi:10.1093/nar/gkw972.

A. Waterhouse, M. Bertoni, S. Bienert, G. Studer, G. Tauriello, R. Gumienny, F. T. Heer, T. A. P. de Beer, C. Rempfer, L. Bordoli, R. Lepore, T. Schwede, SWISS-MODEL: homology modelling of protein structures and complexes. Nucleic Acids Res. 46, W296–W303 (2018).

Z. Wei, Y. Xue, H. Xu, W. Gong, Crystal structure of the C-terminal domain of S.cerevisiae eIF5. J. Mol. Biol. 359, 1–9 (2006).

Y. Yu, A. Marintchev, V. G. Kolupaeva, A. Unbehaun, T. Veryasova, S.-C. Lai, P. Hong, G. Wagner, C. U. T. Hellen, T. V Pestova, Position of eukaryotic translation initiation factor eIF1A on the 40S ribosomal subunit mapped by directed hydroxyl radical probing. Nucleic Acids Res. 37, 5167–82 (2009).

J. Zeman, Y. Itoh, Z. Kukačka, M. Rosůlek, D. Kavan, T. Kouba, M. E. Jansen, M. P. Mohammad, P. Novák, L. S. Valášek, Binding of eIF3 in complex with eIF5 and eIF1 to the 40S ribosomal subunit is accompanied by dramatic structural changes. Nucleic Acids Res. 47, 8282–8300 (2019).

K. Zhang. “Gctf: Real-time CTF determination and correction.” J Struct Biol 193, 1–12 (2016).

S. Q. Zheng, E. Palovcak, J. P. Armache, K. A. Verba, Y. Cheng and D. A. Agard (2017). “MotionCor2: anisotropic correction of beam-induced motion for improved cryo-electron microscopy.” Nat Methods 14, 331–332.

J. Zivanov, T. Nakane, B. O. Forsberg, D. Kimanius, W. J. Hagen, E. Lindahl and S. H. Scheres (2018). “New tools for automated high-resolution cryo-EM structure determination in RELION-3.” Elife 7.

