## Supplementary text and figures for "Structural differences in translation initiation between pathogenic trypanosomatids and their mammalian hosts"

**SUPPLEMENTARY INFORMATION**

***Construction of plasmids***

List of all primers and gene strings used throughout this study is shown below.

pGL4-CMV-h3c was made by inserting the *Pme*I-*Fse*I digested PCR product obtained with primers AH-h3c-PmeI and AH-h3c-FseI using HeLa cDNA as a template into *Pme*I-*Fse*I digested pGL4-CMV.

pGL4-CMV-h3a was made by inserting the *Pme*I-*Fse*I digested PCR product obtained with primers AH-h3a-PmeI and AH-h3a-FseI using YCpLV018 as a template into *Pme*I-*Fse*I digested pGL4-CMV.

pGL4-CMV-h3m was made by inserting the *Eco*RI-*Fse*I digested PCR product obtained with primers AH-h3m-EcoRI and AH-h3m-FseI using HeLa cDNA as a template into *Eco*RI-*Fse*I digested pGL4-CMV.

pGL4-CMV-h3k was made by inserting the *Eco*RI-*Fse*I digested PCR product obtained with primers AH-h3k-EcoRI and AH-h3k-FseI using pFASTBAC1-eIF3k as a template into *Eco*RI-*Fse*I digested pGL4-CMV.

pGL4-CMV-h3d was made by inserting the *Eco*RI-*Fse*I digested PCR product obtained with primers AH-h3d-EcoRI and AH-h3d-FseI using pFASTBAC1-eIF3d as a template into *Eco*RI-*Fse*I digested pGL4-CMV.

pGL4-CMV-h3e was made by inserting the *Eco*RI-*Fse*I digested PCR product obtained with primers AH-h3e-EcoRI and AH-h3e-FseI using pFASTBAC1-eIF3e as a template into *Eco*RI-*Fse*I digested pGL4-CMV.

pGEX-heIF1 was made by inserting the *Bam*HI-*Sal*I digested PCR product obtained with primers DS-eIF1-BamHI and DS-eIF1-SalI using HeLa cDNA as a template into *Bam*HI-*Sal*I digested pGEX-5X-3.

pGEX-heIF5 was made by inserting the *Eco*RI-*Sal*I digested PCR product obtained with primers SW-heIF5-EcoRI and SW-heIF5-SalI-R using HeLa cDNA as a template into *Eco*RI-*Sal*I digested pGEX-5X-3.

pGEX-heIF2β was made by inserting the *Bam*HI-*Sal*I digested PCR product obtained with primers DS-eIF2β-BamHI and DS-eIF2β-SalI using HeLa cDNA as a template into *Bam*HI-*Sal*I digested pGEX-5X-3.

pGL4-CMV-h3c-1-325; pGL4-CMV-h3c-326-913; pGL4-CMV-h3c-30-325 and pGL4-CMV-h3c-130-325 was made by inserting the *Pme*I-*Fse*I digested PCR product obtained with primers AH-h3c-PmeI and AH-h3c-325-FseI; TS-h3c-326-PmeI and AH-h3c-FseI; TS-h3c-30-325-PmeI and AH-h3c-325-FseI; TP-h3c-130-325-PmeI and AH-h3c-325-FseI; respectively, using pGL4-CMV-h3c as a template into *Pme*I-*Fse*I digested pGL4-CMV-h3c.

pGL4-CMV-h3c-1-325-d171-240 was made by inserting the *Pme*I-*Fse*I digested gene string pGL4-CMV-h3c-1-325-d171-240 (GeneArt™ Strings™ DNA Fragments, Invitrogen) into *Pme*I-*Fse*I digested pGL4-CMV.

pGL4-CMV-eIF2β and pGL4-CMV-eIF2β-1-309 was made by inserting the *Eco*RI-*Fse*I digested PCR product obtained with primers TP pGL4-CMV-heIF2β-EcoRI and TP-pGL4-CMV-eIF2β-FseI; TP-pGL4-CMV-heIF2β-EcoRI and TP-pGL4-CMV-eIF2β-1-309-FseI; respectively, using pGEX-heIF2β as a template into *Eco*RI-*Fse*I digested pGL4-CMV.

pGEX-heIF1-box-Ala-102-113 was made by inserting the *Bam*HI-*Sal*I digested gene string heIF1-box-Ala-102-113 (GeneArt™ Strings™ DNA Fragments, Invitrogen) into *Bam*HI-*Sal*I digested pGEX-heIF1.

pGEX-teIF1 and pGEX-teIF5 was made by inserting the *Bam*HI-*Sal*I digested PCR product obtained with primers TP-pGEX-5X3-teIF1-BamHI and TP-pGEX-5X3-teIF1-SalI; TP-pGEX-5X3-teIF5-BamHI and TP-pGEX-5X3-teIF5-SalI; respectively, using *T.cruzi* genomic DNA as a template into *Bam*HI-*Sal*I digested pGEX-5X-3.

pGL4-CMV-teIF5 was made by inserting *Eco*RI-*Fse*I digested PCR product obtained with primers TP-pGL4-teIF5-EcoRI and TP-pGL4-teIF5-FseI using pGEX-teIF5 as a template into *Eco*RI-*Fse*I digested pGL4-CMV.

pGL4-CMV-teIF3c was made by inserting the *Pme*I-*Fse*I digested PCR product obtained with primers TP-pGL4-CMV-teIF3c-PmeI and TP-pGL4-CMV-teIF3c-FseI using *T.cruzi* genomic DNA as a template into *Pme*I-*Fse*I digested pGL4-CMV.

pGL4-CMV-teIF3c-1-172; pGL4-CMV-teIF3c-14-172; pGL4-CMV-teIF3c-39-172 was made by inserting the *Pme*I-*Fse*I digested PCR product obtained with primers TP-pGL4-CMV-teIF3c-PmeI and TPpGL4-CMV-teIF3c-172-FseI; TP-pGL4-CMV-teIF3c-14-PmeI and TPpGL4-CMV-teIF3c-172-FseI; TP-pGL4-CMV-teIF3c-39-PmeI and TP-pGL4-CMV-teIF3c-172-FseI; respectively, using pGL4-CMV-teIF3c as a template into *Pme*I-*Fse*I digested pGL4-CMV.

pGEX-teIF3c-1-172 was made by inserting *Bam*HI-*Eco*RI digested PCR product obtained with primers TP-pGEX-teIF3c-BamHI and TP-pGEX-teIF3c-1-172-EcoRI using pGL4-CMV-teIF3c-1-172 as a template into *Bam*HI-*Eco*RI digested pGEX-5X-3.

pGEX-heIF3e was made by inserting the *Bam*HI-*Sal*I digested PCR product obtained with primers TP-pGEX-5X3-eIF3e-BamHI and TP-pGEX-5X3-eIF3e-SalI using pGL4-CMV-h3e as a template into *Bam*HI-*Sal*I digested pGEX-5X-3.

pGEX-heIF3e-del-244-252 and pGEX-heIF3e-I246A-Q247A-T248A was made by inserting the *Bam*HI-*Bgl*II digested gene string pGEX-heIF3e-delta-244-252 and gene string pGEX-heIF3e-I246A-Q247A-T248A (GeneArt™ Strings™ DNA Fragments, Invitrogen) respectively, into *Bam*HI-*Bgl*II digested pGEX-5X3-heIF3e.

pGL4-CMV-h3d-W16A-G17A-P18A was made by inserting the *Eco*RI-*Pvu*II digested gene string pGL4-CMV-h3d-W16A-G17A-P18A (GeneArt™ Strings™ DNA Fragments, Invitrogen) into *Eco*RI-*Pvu*II digested pGL4-CMV-h3d.
pGL4-CMV-h3d-19-548; pGL4-CMV-h3d-1-114; pGL4-CMV-h3d-19-114 was made by inserting the *Eco*RI-*Fse*I digested PCR product obtained with primers TP-pGL4-CMV-h3d-19-EcoRI and AH-h3d-FseI; AH-h3d-EcoRI and TP-pGL4-CMV-h3d-114-FseI; TP-pGL4-CMV-h3d-19-EcoRI and TP-pGL4-CMV-h3d-114-FseI; respectively, using pGL4-CMV-h3d as a template into *Eco*RI-*Fse*I digested pGL4-CMV.
pGEX-heIF3d was made by inserting the *Bam*HI-*Sal*I digested PCR product obtained with primers TP-pGEX-heIF3d-BamHI and TP-pGEX-heIF3d-SalI using pGL4-CMV-h3d as a template into *Bam*HI-*Sal*I digested pGEX-5X-3.
pGEX-heIF3c was made by inserting the *Eco*RI-*Sal*I digested PCR product obtained with primers TP-pGEX-eIF3c-EcoRI and TP-pGEX-eIF3c-SalI using pGL4-CMV-h3c as a template into *Eco*RI-*Sal*I digested pGEX-5X-3.
pGEX-heIF3a was made by inserting the *Sal*I-*Not*I digested PCR product obtained with primers TP-pGEX-eIF3a-SalI and TP-pGEX-eIF3a-NotI using pGL4-CMV-h3a as a template into *Sal*I-*Not*I digested pGEX-5X-3.
pEX-teIF3c-1-172-GST was made by inserting the *Asi*SI-*Mlu*I digested PCR product obtained with primers TP-teIF3c-AsiSI and TP-teIF3c-1-172-MluI using pGL4-CMV-teIF3c-1-172 as a template into *Bse*RI-*Mlu*I digested pEX-C-GST (OriGene; PS100083)
pEX-teIF5-GST was made by inserting the *Asi*SI-*Mlu*I digested PCR product obtained with primers TP-teIF5-AsiSI and TP-teIF5-MluI using pGEX-teIF5 as a template into *Bse*RI-*Mlu*I digested pEX-C-GST (OriGene; PS100083)

**List of published plasmids used in this study:**
YCpLV018 (Valášek et al., 1998)
pGEX-5X-3 (Smith et al., 2014)
pGL4-CMV (Wagner et al., 2014)
pFASTBAC1-eIF3k (Fraser et al., 2004)
pFASTBAC1-eIF3d (Fraser et al., 2004)
pFASTBAC1-eIF3e (Fraser et al., 2004)

**List of primers**

| Primer | Sequence |
|---|---|
| AH-h3c-PmeI | ATATAGTTTAAACGCCATGTCGCGGTTTTTCACC |
| AH-h3c-FseI | ATATAGGCCGGCCTCAGTAGGCCGTCTGAGACTG |
| AH-h3a-PmeI | ATATAGTTTAAACAAGATGCCGGCCTATTTTCAG |
| AH-h3a-FseI | ATATAGGCCGGCCTTAACGTCGTACTGTGGTCCA |
| AH-h3m-EcoRI | ATATAGAATTCACCATGAGCGTCCCGGCCTTC |
| AH-h3m-FseI | ATATAGGCCGGCCTCAGGTATCAGAAAGACTCAA |
| AH-h3k-EcoRI | ATATAGAATTCGTCATGGCGATGTTTGAGCAG |
| AH-h3k-FseI | ATATAGGCCGGCCTTACTGGGAGGAGGCCATGAT |
| AH-h3d-EcoRI | ATATAGAATTCAAGATGGCAAAGTTCATGACA |
| AH-h3d-FseI | ATATAGGCCGGCCTTAAGTTTCTTCCTCTTCTTCTTCCTC |
| AH-h3e-EcoRI | ATATAGAATTCAAGATGGCGGAGTACGACTTG |
| AH-h3e-FseI | ATATAGGCCGGCCTCAGTAGAAGCCAGAATCTTG |
| DS-eIF1-BamHI | ATCGGATCCATATGTCCGCTATCCAGAACC |
| DS-eIF1-SalI | TGTGTCGACTTAAAACCCATGAACCTTCAG |
| SW-heIF5-EcoRI | ATAGAATTCGATGTCTGTCAATGTCAACC |
| SW-heIF5-SalI-R | ACTAGTCGACTTAAATGGCATCAATATCG |
| DS-eIF2β-BamHI | ATCGGATCCATATGTCTGGGGACGAGATG |
| DS-eIF2β-SalI | CGTGTCGACTTAGTTAGCTTTGGCACG |
| AH-h3c-325-FseI | ATATACGGCCGGCCTCAGGTGATCTCAGTTCCCTTGGC |

| | |
|---|---|
| TS-h3c-326-PmeI | ATATAGTTTAAACGCCATGCATGCTGTTGTTATCAAGAAA CTG |
| TS-h3c-30-325-PmeI | ATATAGTTTAAACGCCATGAACTATGGCAAACAGCCATTG |
| TP-h3c-130-325-PmeI | GCCGCGTTTAAACGCCATGAACAAGAACAATGCCAAGGC |
| TP-pGL4-CMV-heIF2β-EcoRI | CGCCAGAATTCACCATGTCTGGGGACGAGATGATT |
| TP-pGL4-CMV-eIF2β-FseI | ATATAGGCCGGCCTTAGTTAGCTTTGGCACGGAG |
| TP-pGL4-CMV-eIF2β-1-309-FseI | CGCCAGGCCGGCCTTAACATCTAGAATGACAAGTTTC |
| TP-pGEX-5X3-teIF1-BamHI | ACCTAGGATCCCCATGCTAAACAACGAGCTCGCTAACC |
| TP-pGEX-5X3-teIF1-SalI | ACCGAGTCGACTTAGTTCAGAGAGTGGATCTC |
| TP-pGEX-5X3-teIF5-BamHI | ACCGAGGATCCCCATGTCGGTTCCAATGATACCCATTG |
| TP-pGEX-5X3-teIF5-SalI | CGCTAGTCGACCTATGTCGATCCTACAAGCCATTCGAC |
| TP-pGL4-teIF5-EcoRI | GCTGTGAATTCGCCACCATGTCGGTTCCAATGATACCC |
| TP-pGL4-teIF5-FseI | TATATGGCCGGCCCTATGTCGATCCTACAAGCCATTC |
| TP-pGL4-CMV-teIF3c-PmeI | CGCCGGTTTAAACACCATGAGCAACTTTTTGATGTCAGC GACAGTG |
| TP-pGL4-CMV-teIF3c-FseI | ATATAGGCCGGCCGTTAAAATCCTCCTCTACCACGTCCTC GAC |
| TP-pGL4-CMV-teIF3c-14-PmeI | CGCGCGTTTAAACACCATGCTGGATGAGGTCATACATCAC GATG |
| TP-pGL4-CMV-teIF3c-39-PmeI | AGCCAGTTTAAACACCATGACCGATGATGAGGACGCGGA TG |
| TP-pGL4-CMV-teIF3c-172-FseI | TATATGGCCGGCCGTTACTCCTCACCCTGTCCTTCATC |
| TP-pGEX-teIF3c-BamHI | GCGGCGGATCCCCATGAGCAACTTTTTGATGTC |
| TP-pGEX-teIF3c-1-172-EcoRI | GCCGCGAATTCCTTTACTCCTCACCCTGTCCTTC |
| TP-pGEX-5X3-eIF3e-BamHI | ATATAGGATCCCCATGGCGGAGTACGACTTGAC |
| TP-pGEX-5X3-eIF3e-SalI | ATGCCGTCGACTCAGTAGAAGCCAGAATCTTG |
| TP-pGL4-CMV-h3d-19-EcoRI | CTGCAGAATTCAAGATGTGTGCGGTTCCCGAGCAG |
| AH-h3d-FseI | ATATAGGCCGGCCTTAAGTTTCTTCCTCTTCTTCTTCCTC |
| AH-h3d-EcoRI | ATATAGAATTCAAGATGGCAAAGTTCATGACA |
| TP-pGL4-CMV-h3d-114-FseI | ACGTAGGCCGGCCTTACATGTTCCGACGATCTTTGTC |
| TP-pGEX-heIF3d-BamHI | ATATCGGATCCCCATGGCAAAGTTCATGACACCC |
| TP-pGEX-heIF3d-SalI | GCGCGGTCGACTTAAGTTTCTTCCTCTTCTTCTTCCTC |
| TP-pGEX-eIF3c-EcoRI | ACCGAGAATTCCATGTCGCGGTTTTTCACCACC |
| TP-pGEX-eIF3c-SalI | TATATGTCGACTCAGTAGGCCGTCTGAGACTG |
| TP-pGEX-eIF3a-SalI | ACCTAGTCGACATGCCGGCCTATTTTCAGAGG |
| TP-pGEX-eIF3a-NotI | ATATAGCGGCCGCTTAACGTCGTACTGTGGTCCA |
| TP-teIF3c-AsiSI | GCACCGCGATCGCATGAGCAACTTTTTGATGTCAGCG |
| TP-teIF3c-1-172-MluI | ATATAACGCGTCTCCTCACCCTGTCCTTCATC |
| TP-teIF5-AsiSI | ACCGAGCGATCGCATGTCGGTTCCAATGATACCC |
| TP-teIF5-MluI | TCGATACGCGTTGTCGATCCTACAAGCCATTC |

**List of gene-strings:**

heIF1-box-Ala-102-113

GGCGACCATCCTCCAAAATCGGATCTGATCGAAGGTCGTGGGATCCATATGTCCGCTATCCAGAA

CCTCCACTCTTTCGACCCCTTTGCTGATGCAAGTAAGGGTGATGACCTGCTTCCTGCTGGCACTG
AGGATTATATCCATATAAGAATTCAACAGAGAAACGGCAGGAAGACCCTTACTACTGTCCAAGG
G
ATCGCTGATGATTACGATAAAAAGAAACTAGTGAAGGCGTTTAAGAAAAAGTTTGCCTGCAATG
G

TACTGTAATTGAGCATCCGGAATATGGAGAAGTAATTCAGCTACAGGGTGACCAACGCAAGAAC
A
TATGCCAGTTCCTCGTAGAGATTGGAGCAGCAGCAGCTGCAGCAGCCGCAGCCGCGGCAGCATA
A
GTCGACTCGAGCGGCCGCATCGTGACTGACTGACGATCTGCCTCGC

pGEX-heIF3e-delta-244-252
GACCATCCTCCAAAATCGGATCTGATCGAAGGTCGTGGGATCCCCATGGCGGAGTACGAC
TTGACTACTCGCATCGCGCACTTTTTGGATCGGCATCTAGTCTTTCCGCTTCTTGAATTT
CTCTCTGTAAAGGAGATATATAATGAAAAGGAATTATTACAAGGTAAATTGGACCTTCTT
AGTGATACCAACATGGTAGACTTTGCTATGGATGTATACAAAAACCTTTATTCTGATGAT
ATTCCTCATGCTTTGAGAGAGAAAAGAACCACAGTGGTTGCACAACTGAAACAGCTTCAG
GCAGAAACAGAACCAATTGTGAAGATGTTTGAAGATCCAGAAACTACAAGGCAAATGCAG
TCAACCAGGGATGGTAGGATGCTCTTTGACTACCTGGCGGACAAGCATGGTTTTAGGCAG
GAATATTTAGATACACTCTACAGATATGCAAAATTCCAGTACGAATGTGGGAATTACTCA
GGAGCAGCAGAATATCTTTATTTTTTTAGAGTGCTGGTTCCAGCAACAGATAGAAATGCT
TTAAGTTCACTCTGGGGAAAGCTGGCCTCTGAAATCTTAATGCAGAATTGGGATGCAGCC
ATGGAAGACCTTACACGGTTAAAAGAGACCATAGATAATAATTCTGTGAGTTCTCCACTT
CAGTCTCTTCAGCAGAGAACATGGCTCATTCACTGGTCTCTGTTTGTTTTCTTCAATCAC
CCCAAAGGTCGCGATAATATTATTGACCTCTTCCTTTATCAGCCACAATATCTTATTCTT
CGCTATTTGACTACAGCAGTCATAACAAACAAGGATGTTCGAAAACGTCGGCAGGTTCTA
AAAGATCTAGTTAAAGTTATTCAACAGGAGTCTTACACATATAAA

pGEX-heIF3e-I246A-Q247A-T248A
GACCATCCTCCAAAATCGGATCTGATCGAAGGTCGTGGGATCCCCATGGCGGAGTACGAC
TTGACTACTCGCATCGCGCACTTTTTGGATCGGCATCTAGTCTTTCCGCTTCTTGAATTT
CTCTCTGTAAAGGAGATATATAATGAAAAGGAATTATTACAAGGTAAATTGGACCTTCTT
AGTGATACCAACATGGTAGACTTTGCTATGGATGTATACAAAAACCTTTATTCTGATGAT
ATTCCTCATGCTTTGAGAGAGAAAAGAACCACAGTGGTTGCACAACTGAAACAGCTTCAG
GCAGAAACAGAACCAATTGTGAAGATGTTTGAAGATCCAGAAACTACAAGGCAAATGCAG
TCAACCAGGGATGGTAGGATGCTCTTTGACTACCTGGCGGACAAGCATGGTTTTAGGCAG
GAATATTTAGATACACTCTACAGATATGCAAAATTCCAGTACGAATGTGGGAATTACTCA
GGAGCAGCAGAATATCTTTATTTTTTTAGAGTGCTGGTTCCAGCAACAGATAGAAATGCT
TTAAGTTCACTCTGGGGAAAGCTGGCCTCTGAAATCTTAATGCAGAATTGGGATGCAGCC
ATGGAAGACCTTACACGGTTAAAAGAGACCATAGATAATAATTCTGTGAGTTCTCCACTT
CAGTCTCTTCAGCAGAGAACATGGCTCATTCACTGGTCTCTGTTTGTTTTCTTCAATCAC
CCCAAAGGTCGCGATAATATTATTGACCTCTTCCTTTATCAGCCACAATATCTTAATGCA
GCTGCGGCAATGTGTCCACACATTCTTCGCTATTTGACTACAGCAGTCATAACAAACAAG
GATGTTCGAAAACGTCGGCAGGTTCTAAAAGATCTAGTTAAAGTTATTCAACAGGAGTCT
TACACATATAAA

pGL4-CMV-h3c-1-325-d171-240
TGGGAGGTCTATATAAGCAGAGCTCTCTGGCTAACTAGAGAACCCACTGCTTACTGGCTTATCGA
AATTAATACGACTCACTATAGGGAGACCCAAGCTGGCTAGCGTTTAAACGCCATGTCGCGGTTTT
TCACCACCGGTTCGGACAGCGAGTCCGAGTCGTCCTTGTCCGGGGAGGAGCTCGTCACCAAACCT
GTCGGAGGCAACTATGGCAAACAGCCATTGTTGCTGAGCGAGGATGAAGAAGATACCAAGAGAG
T
TGTCCGCAGTGCCAAGGACAAGAGGTTTGAGGAGCTGACCAACCTTATCCGGACCATCCGTAATG
CCATGAAGATTCGTGATGTCACCAAGTGCCTGGAAGAGTTTGAGCTCCTGGGAAAAGCATATGGG

AAGGCCAAAAGCATTGTGGACAAAGAAGGTGTCCCCCGGTTCTATATCCGCATCCTGGCTGACCT
AGAGGACTATCTTAATGAGCTTTGGGAAGATAAGGAAGGGAAGAAGAAGATGAACAAGAACAA
TG
CCAAGGCTCTGAGCACCTTGCGTCAGAAGATCCGAAAATACAACCGTGATTTCGAGTCCCATATC
ACAAGCTACAAGCAGAACCCCGAGCAGTCTGCGGATGAAGATGACTCAGAGGAGGAAGAAGGG
AA
ACAAACCGCGCTGGCCTCAAGATTTCTTAAAAAGGCACCCACCACAGATGAGGACAAGAAGGCA
G
CCGAGAAGAAACGGGAGGACAAAGCTAAGAAGAAGCACGACAGGAAATCCAAGCGCCTGGATG
AG
GAGGAGGAGGACAATGAAGGCGGGGAGTGGGAAAGGGTCCGGGGCGGAGTGCCGTTGGTTAAG
GA
GAAGCCAAAAATGTTTGCCAAGGGAACTGAGATCACCTGAGGCCGGCCGCTTCGAGCAGACATG
A
TAAGATACATTGATGAGTTTGGACAAACCACAACTAGAATGCAGTGAAAAAAATGCTTTATTTGT
GAAATTTGTGATGCTATTGCTTTA

pGL4-CMV-h3d-W16A-G17A-P18A

GGCTAGCGTTTAAACGGGCCCTCTAGACTCGAGCGGCCGCCACTGTGCTGGATATCTGCA
GAATTCAAGATGGCAAAGTTCATGACACCCGTGATCCAGGACAACCCCTCAGGCGCGGCT
GCCTGTGCGGTTCCCGAGCAGTTTCGGGATATGCCCTACCAGCCGTTCAGCAAAGGAGAT
CGGCTAGGAAAGGTTGCAGACTGGACAGGAGCCACATACCAAGATAAGAGGTACACAAAT
AAGTACTCCTCTCAGTTTGGTGGTGGAAGTCAATATGCTTATTTCCATGAGGAGGATGAA
AGTAGCTTCCAGCTGGTGGATACAGCGCGCACACAGAAGACGGCCTACCA

***E.Coli* strains used for GST-fusion protein production:**
Rosetta™ 2(DE3) Competent Cells (Merk)
BL21 Star™ (DE3) (Invitrogen)

### SUPPLEMENTARY FIGURES

| | | | | BASIC Spectral Count (# spectra) | |
|---|---|---|---|---|---|
| | | | | BEFORE Gel Filtration | AFTER Gel Filtration |
| Q4E5Z1\|Q4E5Z1_TRYCC | **DDX60** | Uncharacterized protein OS=Trypanosoma cruzi (strain CL Brener) GN=Tc00.1047053508153.1050 | 10 | 263 | 96 |
| Q4DLI2\|Q4DLI2_TRYCC | **ABCE1** | Ribonuclease L inhibitor, putative OS=Trypanosoma cruzi (strain CL Brener) GN=Tc00.10470535086 | 3 | 103 | 31 |
| **40S ribosomal proteins:** | | | | | |
| | | | BASIC Spectral Count (# spectra) | | |
| | | | BEFORE Gel Filtration | | AFTER Gel Filtration |
| **accession** | | **description** | **40S** | **43S** | **43S** |
| Q4D5P4\|Q4D5P4_TRYCC | | 40S ribosomal protein S4 OS=Trypanosoma cruzi (strain CL Brener) GN=Tc00.1047053509683.117 | 131 | 131 | 93 |
| Q4DTN2\|Q4DTN2_TRYCC | | Activated protein kinase C receptor, putative OS=Trypanosoma cruzi (strain CL Brener) GN=Tc00.10 | 100 | 96 | 48 |
| Q4E0Q3\|Q4E0Q3_TRYCC | | 40S ribosomal protein S5, putative OS=Trypanosoma cruzi (strain CL Brener) GN=Tc00.1047053506 | 65 | 51 | 40 |
| Q4DZ41\|RS3A2_TRYCC | | 40S ribosomal protein S3a-2 OS=Trypanosoma cruzi (strain CL Brener) GN=Tc00.1047053511001.9 | 98 | 84 | 46 |
| Q4E093\|Q4E093_TRYCC | | 40S ribosomal protein S18, putative OS=Trypanosoma cruzi (strain CL Brener) GN=Tc00.104705350 | 75 | 73 | 64 |
| Q4DSU0\|Q4DSU0_TRYCC | | 40S ribosomal protein S6 OS=Trypanosoma cruzi (strain CL Brener) GN=Tc00.1047053510769.49 P | 89 | 72 | 58 |
| Q4CLU9\|Q4CLU9_TRYCC | | 40S ribosomal protein S8 OS=Trypanosoma cruzi (strain CL Brener) GN=Tc00.1047053511069.20 P | 66 | 60 | 46 |
| Q4D4L4\|Q4D4L4_TRYCC | | 40S ribosomal protein S11, putative OS=Trypanosoma cruzi (strain CL Brener) GN=Tc00.104705350 | 58 | 47 | 39 |
| Q4D6I5\|Q4D6I5_TRYCC | | 40S ribosomal protein S14, putative OS=Trypanosoma cruzi (strain CL Brener) GN=Tc00.104705340 | 60 | 61 | 37 |
| Q4CUL0\|Q4CUL0_TRYCC | | 40S ribosomal protein S3, putative OS=Trypanosoma cruzi (strain CL Brener) GN=Tc00.1047053430 | 81 | 72 | 37 |
| Q4D6N9\|Q4D6N9_TRYCC | | Ribosomal protein S19, putative OS=Trypanosoma cruzi (strain CL Brener) GN=Tc00.104705350401 | 39 | 36 | 36 |
| Q4DY30\|Q4DY30_TRYCC | KSRP | RNA-binding protein, putative OS=Trypanosoma cruzi (strain CL Brener) GN=Tc00.1047053511727.2 | 79 | 72 | 29 |
| Q4D4S1\|Q4D4S1_TRYCC | | 40S ribosomal protein S9, putative OS=Trypanosoma cruzi (strain CL Brener) GN=Tc00.1047053504 | 38 | 38 | 28 |
| Q4CUC8\|Q4CUC8_TRYCC | | Ribosomal protein S7, putative OS=Trypanosoma cruzi (strain CL Brener) GN=Tc00.1047053506593 | 84 | 80 | 25 |
| Q4CQU0\|Q4CQU0_TRYCC | | 40S ribosomal protein SA OS=Trypanosoma cruzi (strain CL Brener) GN=Tc00.1047053503719.20 P | 65 | 58 | 22 |
| Q4D916\|Q4D916_TRYCC | | 40S ribosomal protein S16, putative OS=Trypanosoma cruzi (strain CL Brener) GN=Tc00.104705350 | 48 | 52 | 19 |
| Q4E0N6\|Q4E0N6_TRYCC | | 40S ribosomal protein S15a, putative OS=Trypanosoma cruzi (strain CL Brener) GN=Tc00.10470535 | 37 | 34 | 15 |
| Q4DIZ9\|Q4DIZ9_TRYCC | | 40S ribosomal protein S2, putative OS=Trypanosoma cruzi (strain CL Brener) GN=Tc00.1047053503 | 80 | 74 | 27 |
| Q4CXN0\|Q4CXN0_TRYCC | | Ubiquitin/ribosomal protein S27a, putative OS=Trypanosoma cruzi (strain CL Brener) GN=Tc00.1047 | 52 | 40 | 14 |
| Q4DTX6\|Q4DTX6_TRYCC | | Ribosomal protein S25, putative OS=Trypanosoma cruzi (strain CL Brener) GN=Tc00.104705350410 | 46 | 44 | 8 |
| Q4DK39\|Q4DK39_TRYCC | | 40S ribosomal protein S17, putative OS=Trypanosoma cruzi (strain CL Brener) GN=Tc00.104705350 | 58 | 57 | 16 |
| Q4E088\|Q4E088_TRYCC | | 40S ribosomal protein S10, putative OS=Trypanosoma cruzi (strain CL Brener) GN=Tc00.104705350 | 52 | 54 | 22 |
| Q4DW69\|Q4DW69_TRYCC | | 40S ribosomal protein S12 OS=Trypanosoma cruzi (strain CL Brener) GN=Tc00.1047053508231.20 | 34 | 39 | 13 |
| Q4CXV6\|Q4CXV6_TRYCC | | 40S ribosomal protein S33, putative OS=Trypanosoma cruzi (strain CL Brener) GN=Tc00.104705350 | 34 | 31 | 17 |
| Q4DTQ1\|Q4DTQ1_TRYCC | | 40S ribosomal protein S23, putative OS=Trypanosoma cruzi (strain CL Brener) GN=Tc00.104705350 | 33 | 28 | 28 |
| Q4D6H7\|Q4D6H7_TRYCC | | Ribosomal protein S20, putative OS=Trypanosoma cruzi (strain CL Brener) GN=Tc00.104705350847 | 34 | 28 | 16 |
| Q4CWD6\|Q4CWD6_TRYCC | | 40S ribosomal protein S13, putative OS=Trypanosoma cruzi (strain CL Brener) GN=Tc00.104705351 | 32 | 30 | 18 |
| Q4DN73\|Q4DN73_TRYCC | | 40S ribosomal protein S27, putative OS=Trypanosoma cruzi (strain CL Brener) GN=Tc00.104705350 | 21 | 17 | 25 |
| Q4DW38\|Q4DW38_TRYCC | | 40S ribosomal protein S24 OS=Trypanosoma cruzi (strain CL Brener) GN=Tc00.1047053507681.150 | 30 | 26 | 15 |
| Q4CMS5\|Q4CMS5_TRYCC | | Ribosomal protein S29, putative OS=Trypanosoma cruzi (strain CL Brener) GN=Tc00.104705351101 | 17 | 15 | 13 |
| Q4DGZ5\|Q4DGZ5_TRYCC | | 40S ribosomal protein S15, putative OS=Trypanosoma cruzi (strain CL Brener) GN=Tc00.104705351 | 23 | 20 | 11 |
| Q4CYE4\|Q4CYE4_TRYCC | | Ribosomal protein S26, putative OS=Trypanosoma cruzi (strain CL Brener) GN=Tc00.104705350380 | 21 | 18 | 12 |
| Q4E3L9\|Q4E3L9_TRYCC | | 40S ribosomal protein S21, putative OS=Trypanosoma cruzi (strain CL Brener) GN=Tc00.104705351 | 24 | 18 | 7 |
| Q4DA48\|Q4DA48_TRYCC | | 40S ribosomal protein S30, putative OS=Trypanosoma cruzi (strain CL Brener) GN=Tc00.104705350 | 2 | 5 | |
| **Initiation factors:** | | | | | |
| | | | BASIC Spectral Count (# spectra) | | |
| | | | BEFORE Gel Filtration | | AFTER Gel Filtration |
| **accession** | | **description** | **40S** | **43S** | **43S** |
| Q4DL69\|Q4DL69_TRYCC | **eIF3a** | Uncharacterized protein OS=Trypanosoma cruzi (strain CL Brener) GN=Tc00.1047053508919.140 P | 86 | 129 | 50 |
| Q4DSL1\|Q4DSL1_TRYCC | **eIF3b** | Translation initiation factor, putative OS=Trypanosoma cruzi (strain CL Brener) GN=Tc00.104705351 | 95 | 159 | 41 |
| Q4E3G1\|Q4E3G1_TRYCC | **eIF3c** | Eukaryotic translation initiation factor 3 subunit 8, putative OS=Trypanosoma cruzi (strain CL Brener) | 63 | 96 | 16 |
| Q4D7F2\|Q4D7F2_TRYCC | **eIF3e** | Eukaryotic translation initiation factor 3 subunit E OS=Trypanosoma cruzi (strain CL Brener) GN=Tc0 | 60 | 103 | 24 |
| Q4E620\|Q4E620_TRYCC | **eIF2 alpha** | Elongation initiation factor 2 alpha subunit, putative OS=Trypanosoma cruzi (strain CL Brener) GN=T | 5 | 105 | 22 |
| Q4DCN0\|Q4DCN0_TRYCC | **eIF3d** | Eukaryotic translation initiation factor 3 subunit 7-like protein, putative OS=Trypanosoma cruzi (strain | 72 | 113 | 16 |
| Q4D452\|Q4D452_TRYCC | **eIF3i** | Eukaryotic translation initiation factor 3 subunit I OS=Trypanosoma cruzi (strain CL Brener) GN=Tc00 | 40 | 69 | 14 |
| Q4D5W3\|Q4D5W3_TRYCC | **eIF3l** | Eukaryotic translation initiation factor 3 subunit L OS=Trypanosoma cruzi (strain CL Brener) GN=Tc0 | 51 | 83 | 25 |
| Q4E3S5\|Q4E3S5_TRYCC | **eIF3h** | Homology with eIF3H (InterPro), Uncharacterized protein OS=Trypanosoma cruzi (strain CL Brener) | 36 | 58 | 8 |
| Q4CUG4\|Q4CUG4_TRYCC | **eIF3g** | Eukaryotic translation initiation factor 3 subunit G OS=Trypanosoma cruzi (strain CL Brener) GN=Tc0 | 43 | 70 | 19 |
| Q4CSE1\|Q4CSE1_TRYCC | **eIF5** | Eukaryotic translation initiation factor 5, putative OS=Trypanosoma cruzi (strain CL Brener) GN=Tc00 | 19 | 115 | 49 |
| Q4DDK1\|Q4DDK1_TRYCC | **eIF3k** | Homology with eIF3K (InterPro), Uncharacterized protein OS=Trypanosoma cruzi (strain CL Brener) | 18 | 29 | 9 |
| Q4DH88\|Q4DH88_TRYCC | **eIF2 beta** | Translation initiation factor, putative OS=Trypanosoma cruzi (strain CL Brener) GN=Tc00.104705350 | 7 | 45 | 25 |
| Q4DQZ2\|Q4DQZ2_TRYCC | **eIF3f** | Uncharacterized protein OS=Trypanosoma cruzi (strain CL Brener) GN=Tc00.1047053510089.200 P | 46 | 68 | 24 |
| Q4CPV7\|Q4CPV7_TRYCC | **eIF2 gamma** | Eukaryotic translation initiation factor 2 subunit, putative OS=Trypanosoma cruzi (strain CL Brener) G | 5 | 62 | 7 |
| Q4CQB1\|Q4CQB1_TRYCC | **eIF1A** | Eukaryotic translation initiation factor 1A, putative (Fragment) OS=Trypanosoma cruzi (strain CL Bre | 4 | 25 | 4 |
| Q4DM75\|Q4DM75_TRYCC | **eIF1** | Protein translation factor SUI1 homolog, putative OS=Trypanosoma cruzi (strain CL Brener) GN=Tc0 | 10 | 18 | 5 |

**Supplementary Figure 1. Mass-spectrometry analysis of the *T. cruzi* 43S PIC.** Composition of the *T. cruzi* 43S PIC in 40S ribosomal proteins and initiation factors. K-DDX60 and ABCE1 were singled out. The analysis compares the 43S related fractions without (labeled 40S) and with GMP-PNP (labeled 43S), before and after Gel-filtration. Accessions, description and spectral counts are indicated for each fraction. Full dataset can be found at the PRIDE partner repository with the dataset identifier PXD016063 (See Methods).

| | Name | | Spectral Count IC |
|---|---|---|---|
| tr\|E9ACL4\|E | DDX60 | Uncharacterized protein OS=Leishmania major GN=LMJF_03_0690 PE=4 SV=1 | 111 |
| tr\|Q4QCE4\| | ABCE1 | Putative ATP-binding cassette protein subfamily E,member 1 OS=Leishmania major GN=ABCE | 101 |

#### 40S ribosomal proteins:

| accession | Name | description | Spectral Count IC |
|---|---|---|---|
| tr\|Q868B1\|C | | 40S ribosomal protein S5 OS=Leishmania major GN=LMJF_11_0960 PE=4 SV=1 | 188 |
| tr\|Q4Q216\|C | | Putative ubiquitin/ribosomal protein S27a OS=Leishmania major GN=LMJF_36_0600 PE=4 SV | 265 |
| tr\|Q4Q1Y2\|C | | Putative 40S ribosomal protein S18 OS=Leishmania major GN=LMJF_36_0940 PE=3 SV=1 | 122 |
| tr\|Q4QG31\| | | 40S ribosomal protein S4 OS=Leishmania major GN=RS4 PE=2 SV=1 | 299 |
| tr\|Q4Q8H1\|C | | 40S ribosomal protein S14 OS=Leishmania major GN=LMJF_28_0960 PE=3 SV=1 | 155 |
| tr\|Q4QC89\|C | | Putative 40S ribosomal protein S23 OS=Leishmania major GN=LMJF_21_1060 PE=3 SV=1 | 90 |
| tr\|Q4Q4A0\|C | | Putative 40S ribosomal protein S3 OS=Leishmania major GN=LMJF_15_0950 PE=4 SV=1 | 99 |
| sp\|P25204\|I | | 40S ribosomal protein S8 OS=Leishmania major GN=RPS8A PE=3 SV=1 | 108 |
| sp\|Q9NE83 | | 40S ribosomal protein S6 OS=Leishmania major GN=RPS6 PE=3 SV=1 | 175 |
| tr\|Q4Q817\|C | | Putative ribosomal protein S29 OS=Leishmania major GN=LMJF_28_2205 PE=4 SV=1 | 62 |
| tr\|Q4Q1V1\|C | | Putative 40S ribosomal protein S9 OS=Leishmania major GN=LMJF_36_1250 PE=2 SV=1 | 98 |
| tr\|Q4Q5P0\|C | | 40S ribosomal protein S2 OS=Leishmania major GN=LMJF_32_0450 PE=3 SV=1 | 144 |
| tr\|Q4Q3M1\| | | Putative 40S ribosomal protein S13 OS=Leishmania major GN=LMJF_19_0390 PE=3 SV=1 | 83 |
| tr\|Q4QH01\| | | Putative 40S ribosomal protein S21 OS=Leishmania major GN=LMJF_11_0760 PE=4 SV=1 | 39 |
| sp\|Q4FX73 | | 40S ribosomal protein S3a OS=Leishmania major GN=LmjF.35.0400 PE=2 SV=1 | 288 |
| tr\|Q4Q8G4\| | | Putative ribosomal protein S20 OS=Leishmania major GN=LMJF_28_1010 PE=3 SV=1 | 99 |
| tr\|Q4Q7P0\|C | | Putative 40S ribosomal protein S30 OS=Leishmania major GN=LMJF_30_0670 PE=4 SV=1 | 36 |
| tr\|Q4QCN7\| | | Putative 40S ribosomal protein S11 OS=Leishmania major GN=LMJF_20_1650 PE=3 SV=1 | 153 |
| sp\|Q4Q0Q0 | | 40S ribosomal protein SA OS=Leishmania major GN=LmjF36.5010 PE=3 SV=1 | 145 |
| tr\|E9AEE8\|E | | 40S ribosomal protein S19-like protein OS=Leishmania major GN=LMJF_29_2860 PE=4 SV=1 | 129 |
| tr\|Q4Q931\|C | | Putative 40S ribosomal protein S33 OS=Leishmania major GN=S33-1 PE=4 SV=1 | 102 |
| tr\|Q4Q1X7\|C | | Putative 40S ribosomal protein S10 OS=Leishmania major GN=LMJF_36_0980 PE=4 SV=1 | 101 |
| tr\|Q4QG97\| | | 40S ribosomal protein S12 OS=Leishmania major GN=LMJF_13_0570 PE=3 SV=1 | 93 |
| tr\|Q4QGW3 | | Putative 40S ribosomal protein S15A OS=Leishmania major GN=LMJF_11_1190 PE=3 SV=1 | 84 |
| tr\|Q4Q9A5\|C | | Putative 40S ribosomal protein S16 OS=Leishmania major GN=LMJF_26_0880 PE=2 SV=1 | 79 |
| tr\|Q4Q806\|C | | Putative 40S ribosomal protein S17 OS=Leishmania major GN=LMJF_28_2555 PE=3 SV=1 | 42 |
| tr\|Q4Q140\|C | | Putative 40S ribosomal protein S27-1 OS=Leishmania major GN=LMJF_36_3750 PE=3 SV=1 | 63 |
| tr\|Q4Q8L6\|C | | Putative ribosomal protein S26 OS=Leishmania major GN=LMJF_28_0540 PE=4 SV=1 | 34 |
| tr\|Q4Q1D2\| | | 40S ribosomal protein S24 OS=Leishmania major GN=S24E-2 PE=3 SV=1 | 120 |
| tr\|Q4Q3G4\| | | Ribosomal protein S25 OS=Leishmania major GN=S25 PE=4 SV=1 | 91 |
| tr\|O43943\|C | RACK1 | LACK OS=Leishmania major PE=4 SV=1 | 58 |
| tr\|Q4Q5K7\|C | KSRP | Putative RNA binding protein OS=Leishmania major GN=LMJF_32_0750 PE=4 SV=1 | 56 |
| tr\|Q4QBV0\| | | Putative 40S ribosomal protein S15 OS=Leishmania major GN=LMJF_22_0420 PE=3 SV=1 | 31 |
| tr\|E9AC32\|E | | Putative ribosomal protein S7 OS=Leishmania major GN=LMJF_01_0410 PE=4 SV=1 | 27 |

#### Initiation factors:

| accession | Name | description | Spectral Count (# spe IC |
|---|---|---|---|
| tr\|Q4QEJ8\|C | eIF3a | Uncharacterized protein OS=Leishmania major GN=LMJF_17_0010 PE=4 SV=1 | 278 |
| tr\|Q4QE62\|C | eIF3b | Putative translation initiation factor OS=Leishmania major GN=LMJF_17_1290 PE=4 SV=1 | 175 |
| tr\|Q4Q6Y6\|C | eIF3d | Eukaryotic translation initiation factor 3 subunit 7-like protein OS=Leishmania major GN=LMJF | 125 |
| tr\|Q4Q833\|C | eIF3e | Eukaryotic translation initiation factor 3 subunit E OS=Leishmania major GN=LMJF_28_2310 F | 84 |
| tr\|Q4Q253\|C | eIF3l | Eukaryotic translation initiation factor 3 subunit L OS=Leishmania major GN=LMJF_36_0250 P | 91 |
| tr\|Q4Q127\|C | eIF3i | Eukaryotic translation initiation factor 3 subunit I OS=Leishmania major GN=LMJF_36_3880 PI | 79 |
| tr\|E9ACP3\|E | eIF2 alpha | Putative elongation initiation factor 2 alpha subunit OS=Leishmania major GN=LMJF_03_0980 | 76 |
| tr\|Q4QIM7\|C | eIF3h | Uncharacterized protein OS=Leishmania major GN=LMJF_07_0640 PE=4 SV=1 | 76 |
| tr\|Q4Q3H3\|C | eIF5 | Putative eukaryotic translation initiation factor 5 OS=Leishmania major GN=LMJF_34_0350 PE | 75 |
| tr\|Q4Q9T0\|C | eIF3f | Uncharacterized protein OS=Leishmania major GN=LMJF_25_1610 PE=4 SV=1 | 67 |
| tr\|Q4Q055\|C | eIF3c | Putative eukaryotic translation initiation factor 3 subunit 8 OS=Leishmania major GN=LMJF_36 | 62 |
| tr\|Q4Q557\|C | eIF3k | Uncharacterized protein OS=Leishmania major GN=LMJF_32_2180 PE=4 SV=1 | 59 |
| tr\|Q4QHR7\| | eIF2 gamma | Putative eukaryotic translation initiation factor 2 subunit OS=Leishmania major GN=LMJF_09_ | 49 |
| tr\|Q4Q2S5\|C | eIF3g | Eukaryotic translation initiation factor 3 subunit G OS=Leishmania major GN=LMJF_34_2700 F | 46 |
| tr\|Q4QAL1\|C | eIF1A | Putative translation factor sui1 OS=Leishmania major GN=LMJF_24_1210 PE=4 SV=1 | 33 |
| tr\|Q4QIB4\|C | eIF2 beta | Translation initiation factor-like protein OS=Leishmania major GN=LMJF_08_0550 PE=4 SV=1 | 29 |
| tr\|Q4QF06\|C | eIF1A | Putative eukaryotic translation initiation factor 1A OS=Leishmania major GN=LMJF_16_0140 F | 27 |

**Supplementary Figure 2. Mass-spectrometry analysis of the *L. Tarentolae* 43S PIC.** Composition of the *L. Tarentolae* 43S PIC in 40S ribosomal proteins and initiation factors. K-DDX60 and ABCE1 were singled out. The analysis of the 43S related fraction was made after supplementation with GMP-PNP (IC), before Gel-filtration. Accessions, description and spectral counts are indicated. Full dataset can be found at the PRIDE partner repository with the dataset identifier PXD016063 (See Methods).

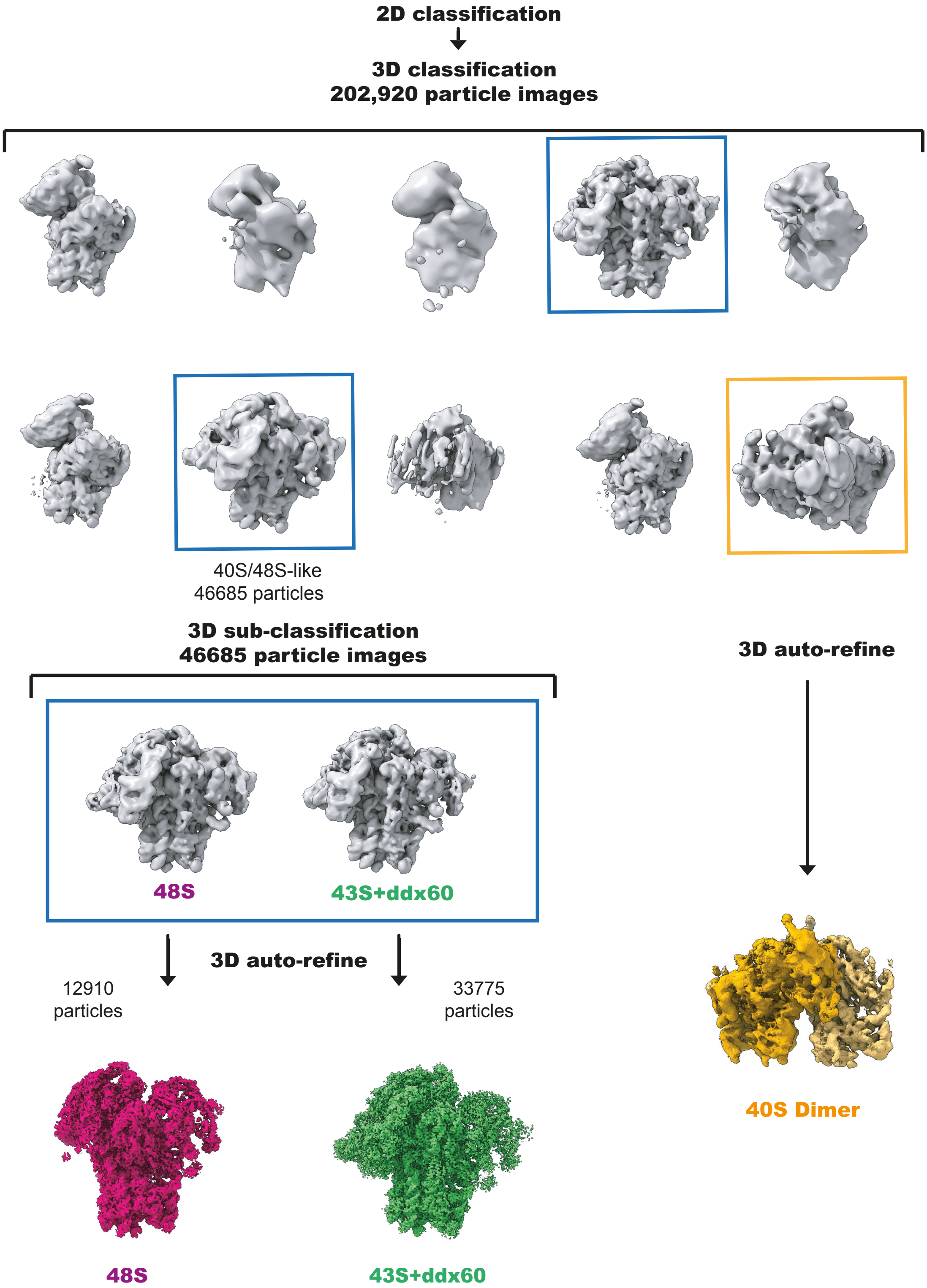

**Supplementary Figure 3. Cryo-EM particle sorting and refinement of the *T. cruzi* 43S PIC.** 2D classification of the 43S PIC particles yielded ~200 000 40S-like particles, after which a run of 3D classification (10 classes) allowed to single out 43S/48S ICs and 40S-dimers. A secondary run of 3D classification allowed the sorting of the 43S PIC particles that generated the main reconstruction analyzed in this study.

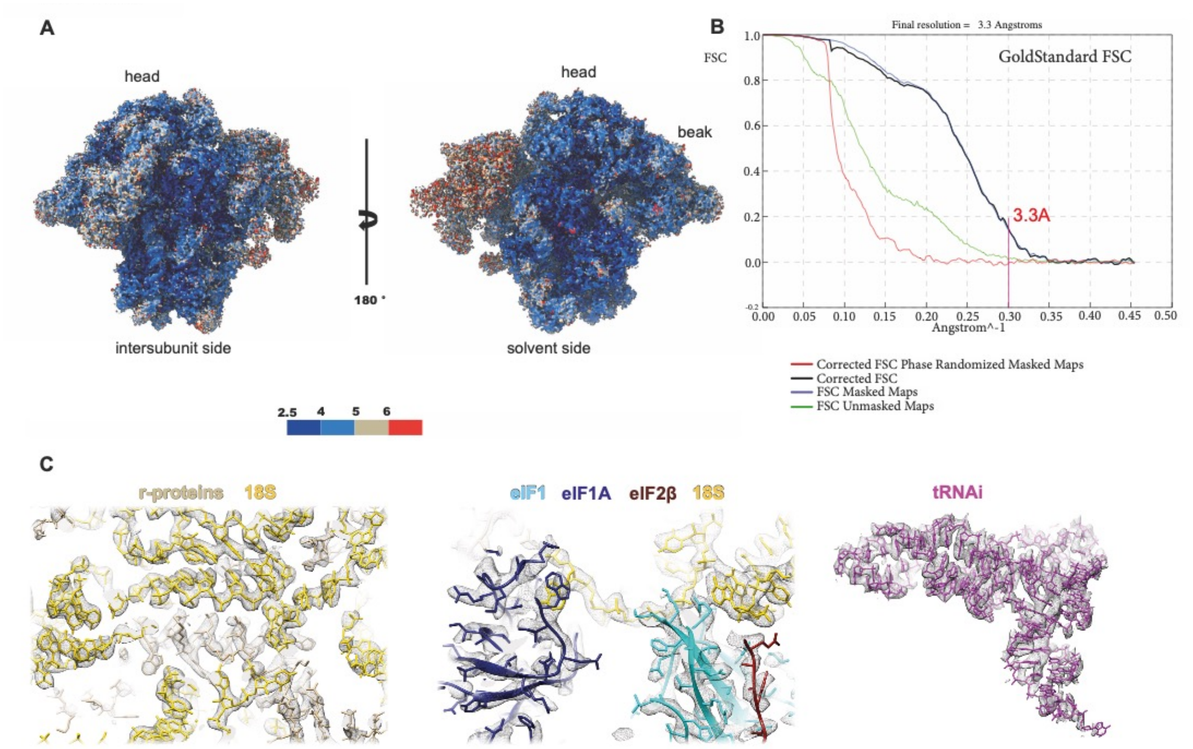

**Supplementary Figure 4. Cryo-EM average and local resolution of the *T. cruzi* 43S PIC.** (**A**) The local resolution varies mainly on eIF3 (ranging from ~3 to ~6 Å), while is varies less on the rest of the structure (ranging from ~2.5 to ~3.5 Å for the 40S, k-DDX60, eIFs 1, 1A and 2b, and from ~3 to ~5 Å for eIFs 2a, 2g and 5). (**B**) The average resolution was measured after applying a soft-edge mask of the 43S PIC shape filtered to 15Å and extended by 3 pixels. (**C**) Blow ups on several features of the complex counting 40S rRNA/r-proteins (left), 18S rRNA interaction with eIF 1 and 1A (middle) and the initiator $tRNA^{Met}$ (right), fitted in their corresponding densities.

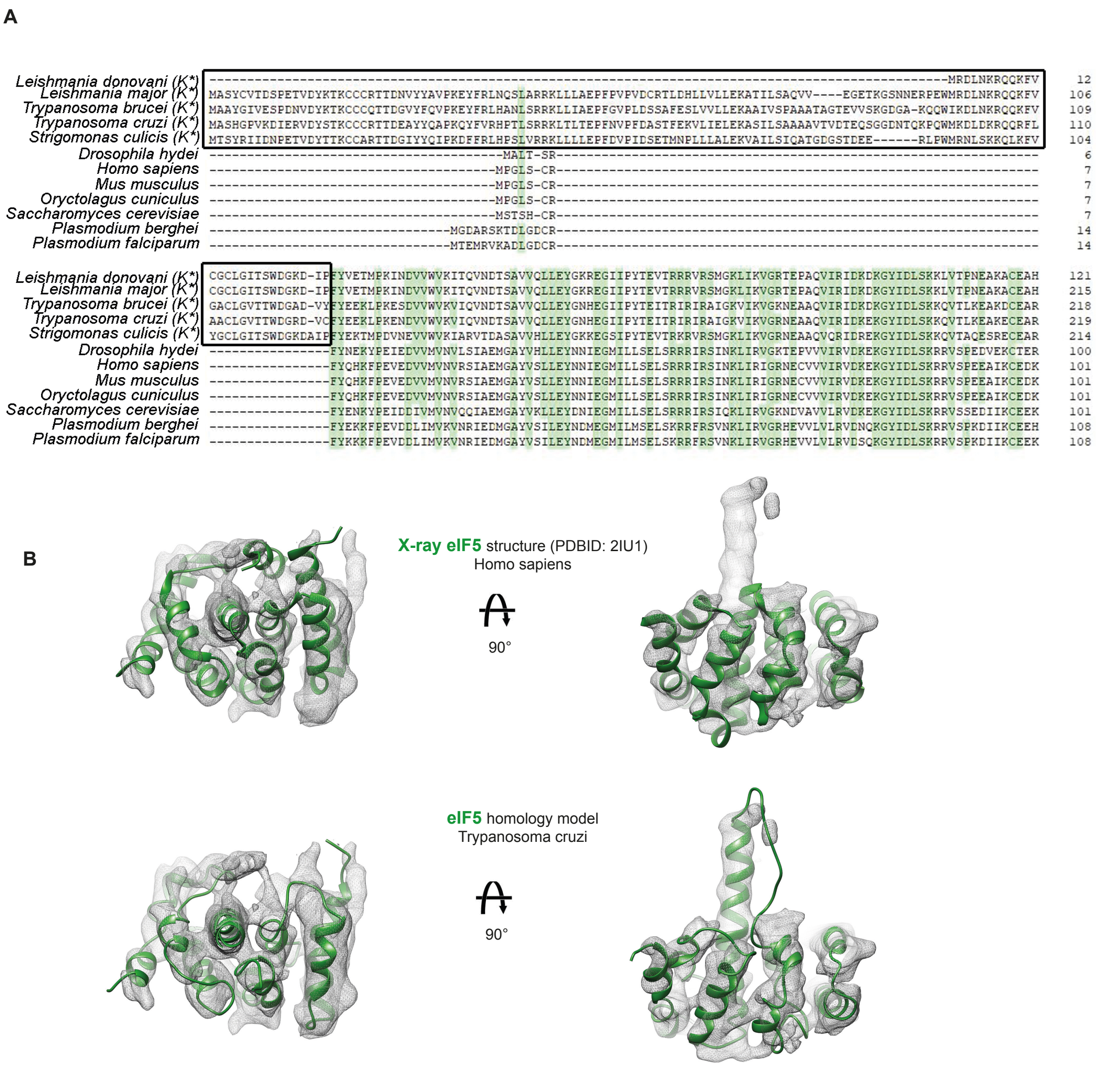

**Supplementary Figure 5. Multiple sequence alignment of the eIF2α NTD among eukaryotes and eIF5 CTD structure.** (**A**) Protein sequence alignment of eIF2α from various eukaryotic organisms was generated by Clone Manger (MultiWay, scoring matrix: Blosum 62). The Kinetoplastida order species are labeled with K*. The kinetoplastidian-specific eIF2α N-terminal domain insertion is marked with a black box. Areas of high matches (60%) are shaded in green. The individual species with the NCBI Reference Sequence numbers or TriTrypDB numbers are as follows: [*Trypanosoma cruzi*] PWV18423.1, [*Trypanosoma brucei*] Tb927.3.2900, [*Leishmania donovani*] AAQ02666.1, [*Leishmania major*] LmjF.03.0980, [*Strigomonas culicis*] EPY26930.1, *[Plasmodium falciparum* NF54] PKC42156.1, [*Plasmodium berghei* ANKA] VUC53995.1, [*Saccharomyces cerevisiae*] ONH75775.1, [*Oryctolagus cuniculus*] XP_002719561.1, [*Mus musculus*] NP_080390.1, [*Drosophila hydei*] XP_023166950.2, [*Homo sapiens*] NP_004085.1. (**B**) Rigid-body fittings of the crystal structure of the human eIF5 CTD in the corresponding *T. cruzi* 43S PIC density (up) and its *T. cruzi* eIF5 CTD homology model in that same density (bottom).

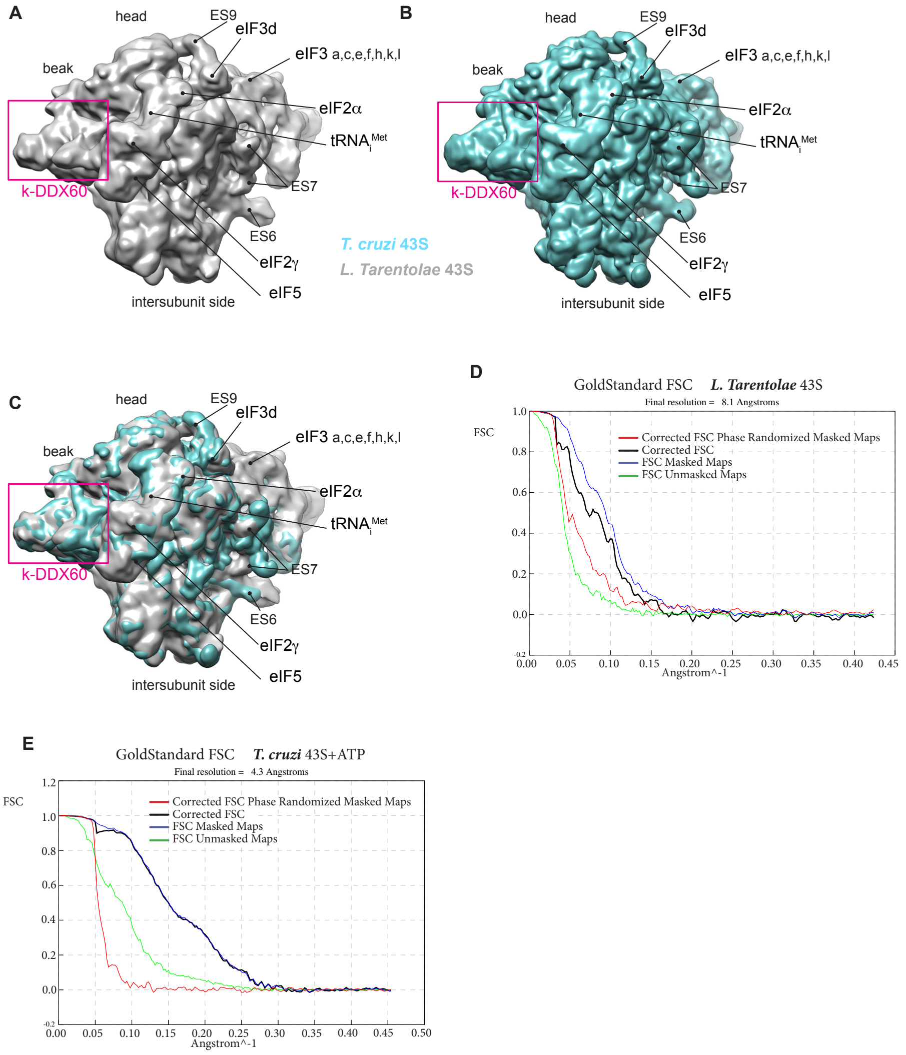

**Supplementary Figure 6. Cryo-EM reconstruction of *L. tarentolae* 43S PIC compared to *T. cruzi* and average resolution.** (**A**) Cryo-EM reconstructions of the *L. tarentolae* 43S PIC. (**B**) Cryo-EM reconstructions of the *T. cruzi* 43S PIC filtered at 8Å. (**C**) Superimposition of (A) and (B). (**D**) Average resolution (8.1Å) of the *L. tarentolae* 43S PIC reconstruction. (**E**) Average resolution (4.3Å) of the cryo-EM reconstruction from the *T. cruzi* 43S complexes supplemented with ATP.

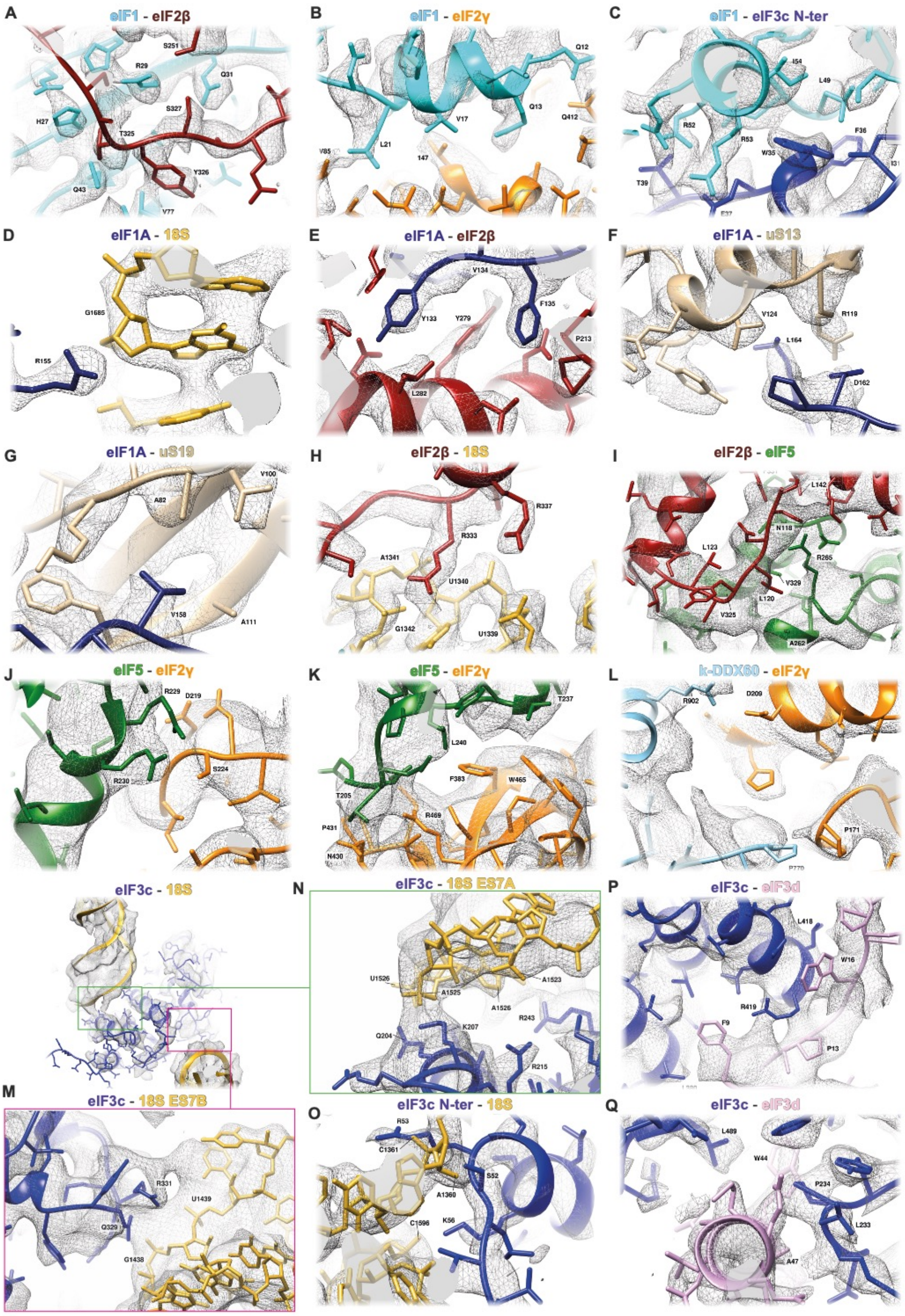

**Supplementary Figure 7. Novel interactions between several eIFs, r-proteins and 18S rRNA, fitted in their corresponding densities.** (**A**) eIF1 with eIF2β. (**B**) eIF1 N-ter tail with eIF2γ. (**C**) eIF1 with eIF3c N-ter. (**D**) eIF1A with the 18S. (**E**) eIF1A with eIF2β. (**F**) eIF1A with uS13. (**G**) eIF1A with uS19. (**H**) eIF2β with the 18S. (**I**) eIF2β with eIF5 CTD. (**J** and **K**) eIF5 CTD with eIF2γ. (**L**) eIF2γ with k-DDX60. (**M**, **N** and **O**) 18S with eIF3c. (**P** and **Q**) eIF3c and eIF3d subunits.

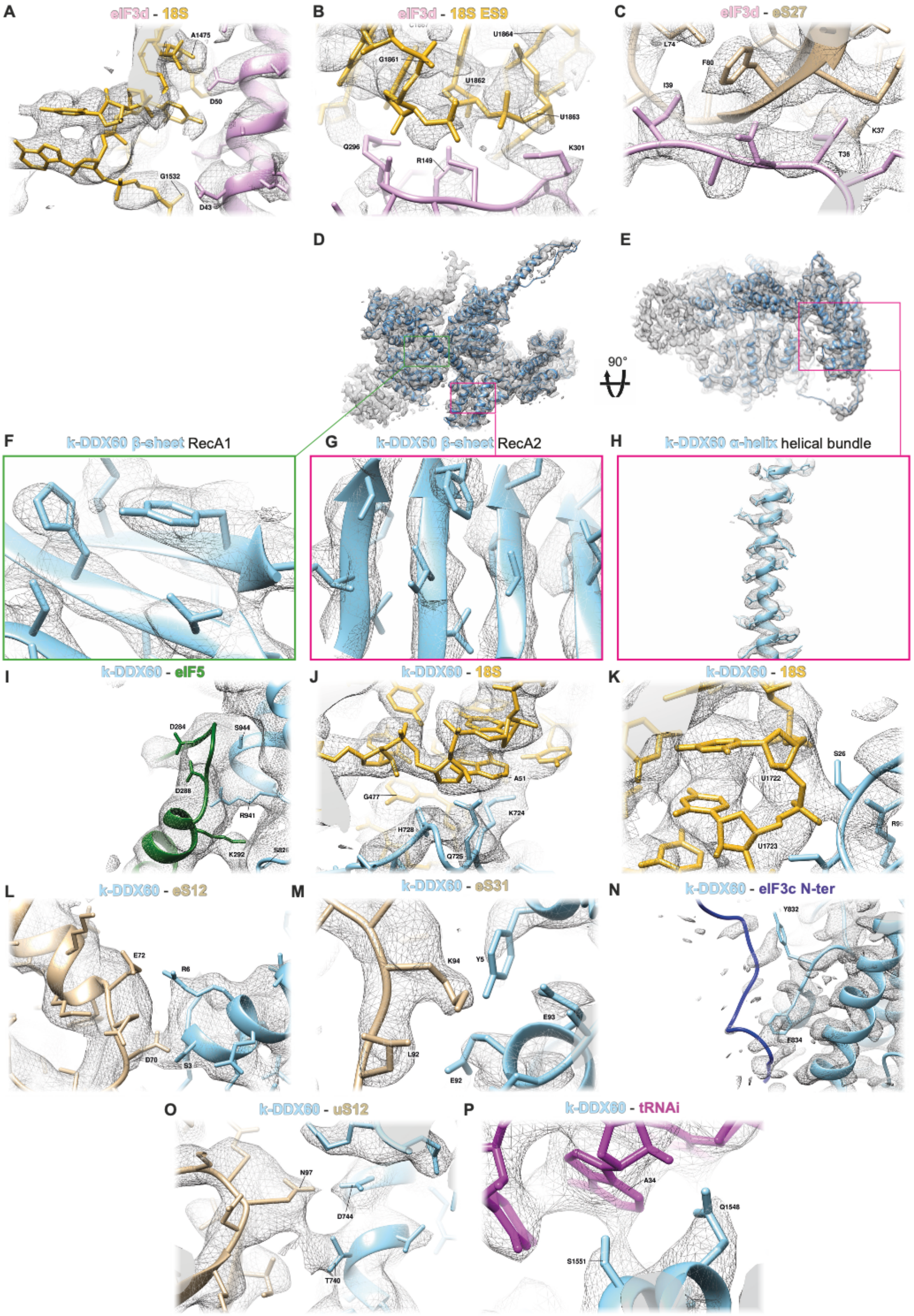

**Supplementary Figure 8. Interactions of eIF3d and k-DDX60 with eIFs, r-proteins and 18S rRNA, fitted in their corresponding densities.** (**A** and **B**) eIF3d with the 18S. (**C**) eIF3d with eS27. (**D** and **E**) k-DDX60 in its corresponding density, viewed from two orientations. (**F**, **G** and **H**) Blow ups on two β-sheets from helicase RecA domains and one buried α-helix from a helical bundle from K-DDX60. Interactions of k-DDX60 with eIF5 (**I**), 18S (**J** and **K**), eS12 (**L**), eS31 (**M**), eIF3c N-ter (**N**), uS12 (**O**) and the initiator $tRNA^{Met}$ (**P**).

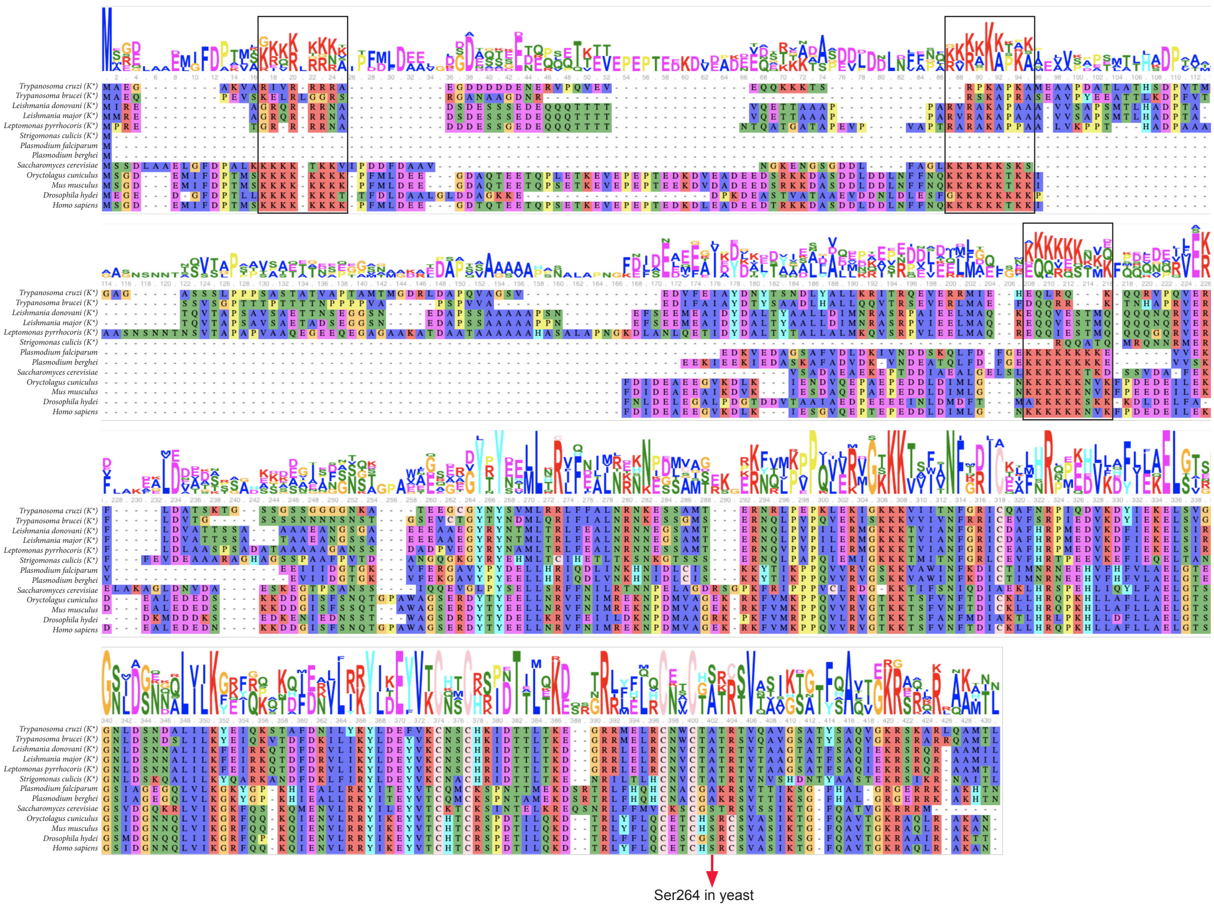

**Supplementary Figure 9. Multiple sequence alignment of eIF2β among eukaryotes.** Protein sequence alignment of eIF2β protein from various eukaryotic organisms. The Kinetoplastida order species are labeled with K*. Consensus is expressed as a sequence logo. The black boxes mark three conserved poly-lysine stretches (dubbed K-boxes) K1, K2 and K3.

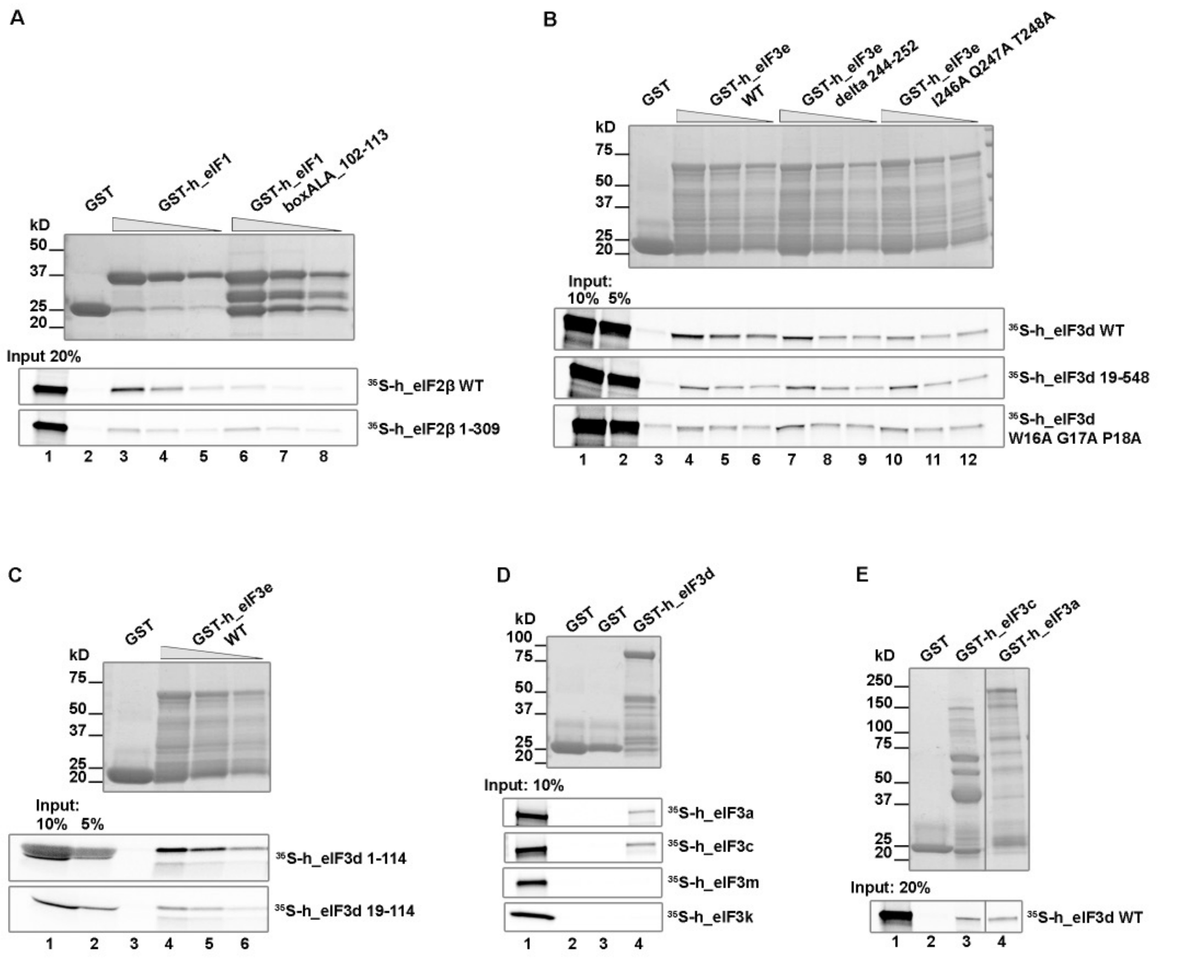

**Supplementary Figure 10. *In vitro* analysis of eIF3 intersubunit interactions.** (**A**) *In vitro* protein-protein binding analysis of the interaction between the *in vitro* translated human $^{35}$S-labeled eIF2β and its C-terminal truncation (eIF2β 1-309) against wild type eIF1 or its mutated variant (eIF1-boxAla-102-113; residues 102-113 substituted with a stretch of alanines) fused to GST. *In vitro* translated proteins were tested for binding with three different dilutions of individual GST-fusion proteins. Lane 1 contains 20% of input amounts of *in vitro*-translated proteins added to each reaction. (**B**) Same as in (A) except that binding between the human wild type eIF3d subunit, its N-terminally truncated form (19-548), and its mutated variant (W16A G17A P18A) against the human wild type eIF3e subunit, or its inner deletion (delta 244-252), or its mutated variant (I246A Q247A T248A) fused to GST was analyzed. Lanes 1 and 2 show 10% and 5% input, respectively. Quantification was performed by the Quantity One software (see Fig. 3J.) (**C**) Same as in (A) except that binding between truncations of the human eIF3d subunit (1-114 and 19-114) and eIF3e fused to GST was analyzed. Quantification is presented in Fig. 3K. (**D**) *In vitro* protein-protein binding analysis of $^{35}$S-labeled eIF3a, eIF3c, eIF3k and eIF3m subunits against eIF3d fused to GST. Lane 1 shows 10% input. (**E**) *In vitro* protein-protein binding analysis of human $^{35}$S-labeled eIF3d against eIF3c and eIF3a subunits fused to GST. Lane 1 shows 20% input.

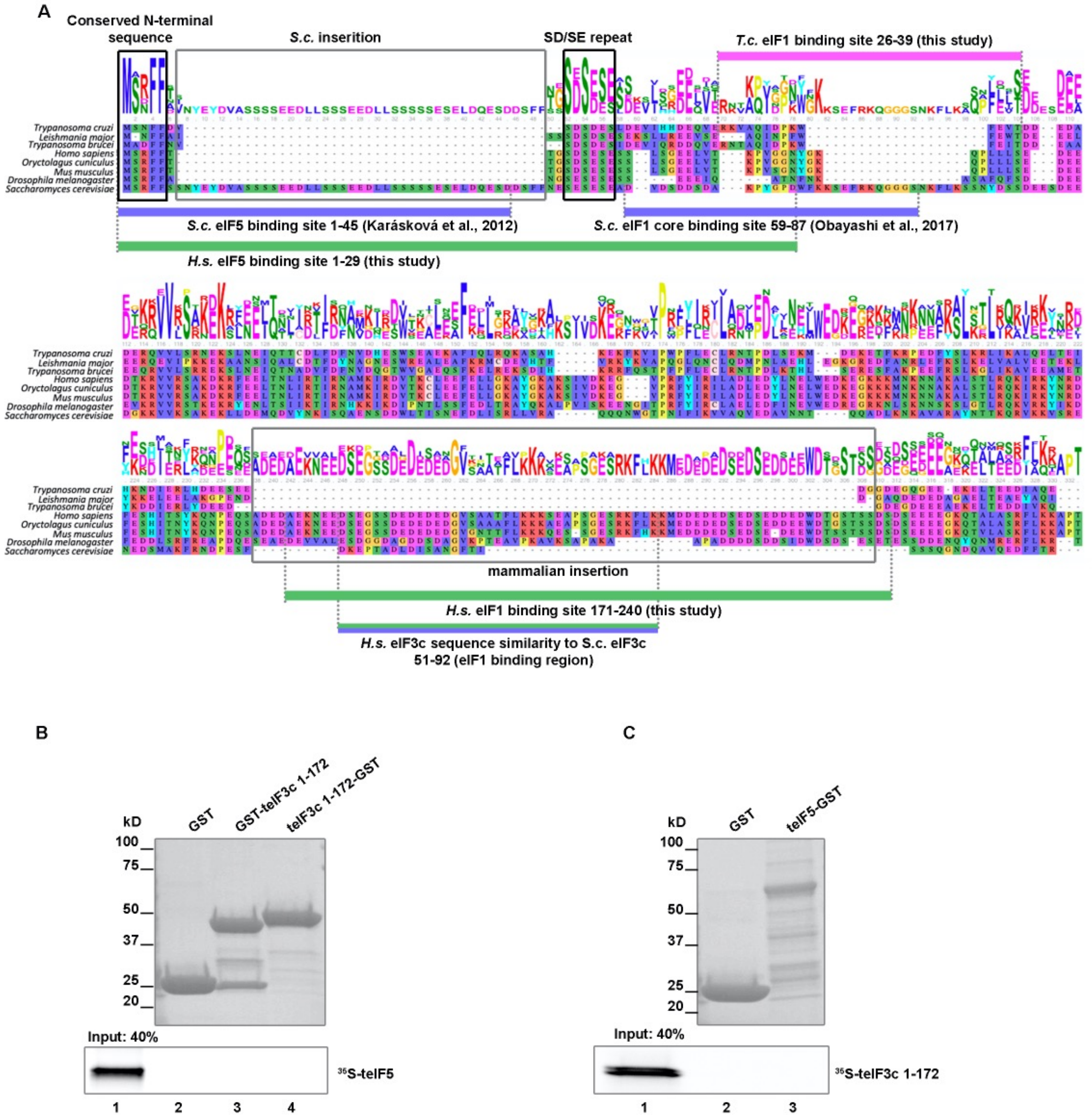

**Supplementary Figure 11. Analysis of eIF1 and eIF5 binding sites on eIF3c.** (**A**) Multiple protein alignment of the N-terminal domain of the eIF3c subunit from indicated species with a consensus expressed as a sequence logo. Specific sequence features mentioned in the main text are boxed. Positions of eIF1- and eIF5-binding sites in the eIF3c-NTD of the selected species identified by us and others are marked by thick lines under or above the alignment; color-coding is as follows: *T.c. – Trypanosoma cruzi* in pink, *S.c. – Saccharomyces cerevisiae* in purple, and *H.s. – Homo sapiens* in green. (**B**) *In vitro* protein-protein binding analysis of the interaction between *T. cruzi* eIF5 and the eIF3c-NTD (residues 1-172) fused with GST either at its N or C terminus. (**C**) Binding analysis of the interaction between *T.cruzi* the eIF3c-NTD (residues 1-172) and eIF5-GST.

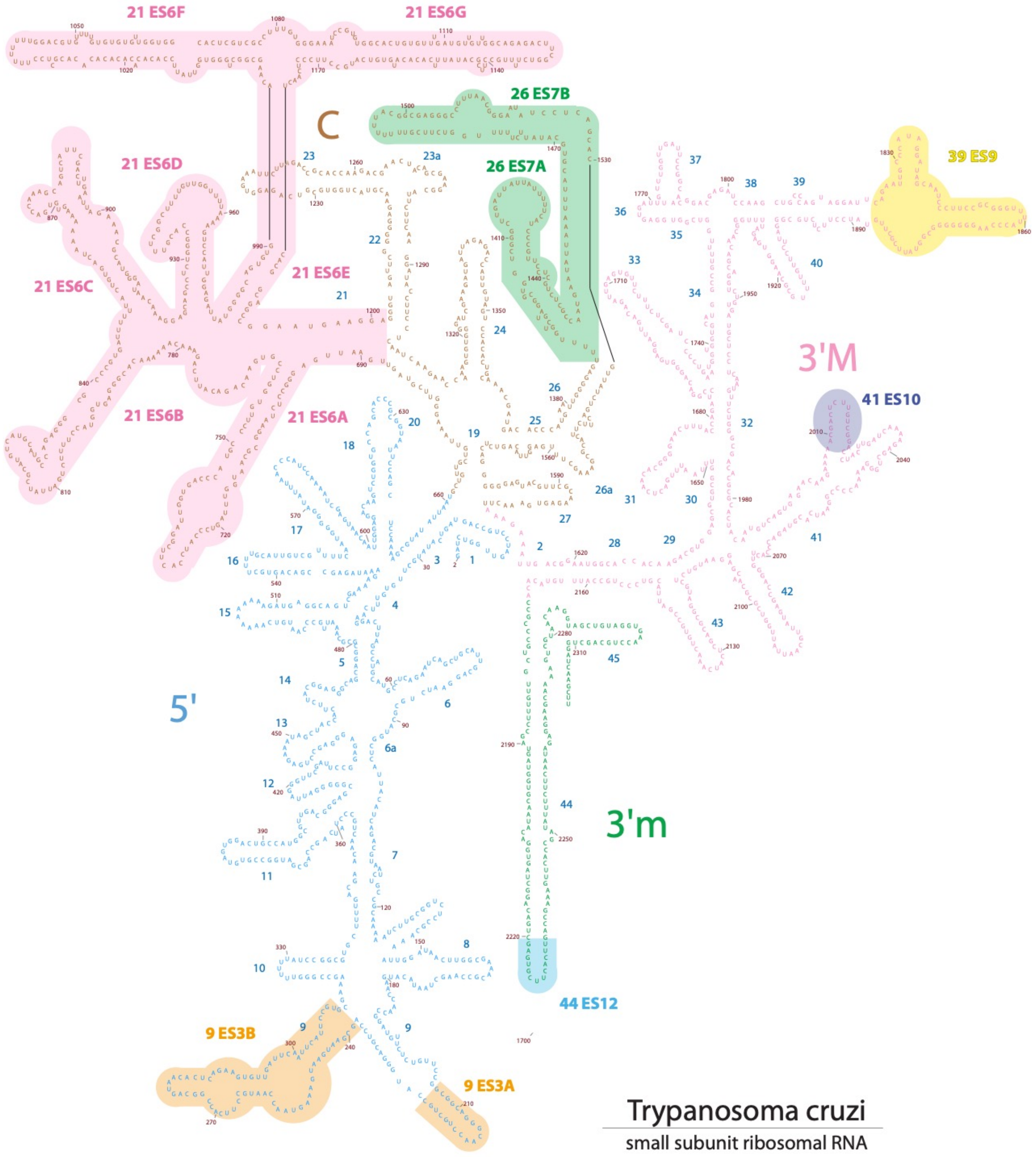

**Figure S12. 2D diagram of the *T. cruzi* 18S structure.** The largest and more relevant expansion segments are highlighted in colored backgrounds.

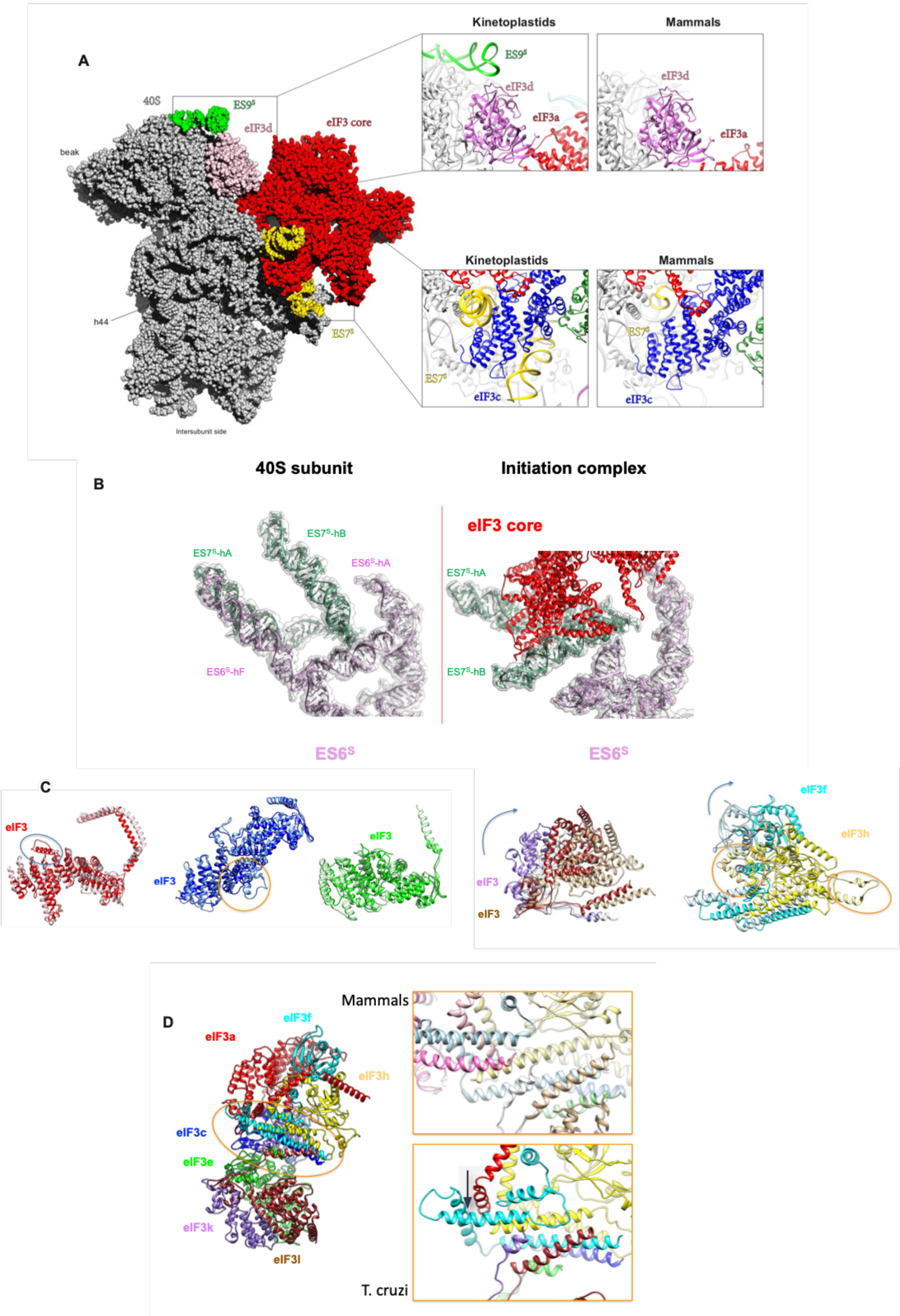

**Supplementary Figure 13. Specific features of Kinetoplastidian eIF3 and its ribosome binding site. (A)** Overall sphere representation of the *T.cruzi* 43S PIC showing kinetoplastidian specific rRNA oversized expansion segments (ESs) in contact with eIF3. Upper panel: comparison of the kinetoplastidian and mammalian eIF3d docking site within the 43S PIC (eIF3d in violet, ES9$^s$ in green, eIF3a in red); lower panel: comparison of the kinetoplastidian and mammalian eIF3c docking site within the 43S PIC (eIF3c in blue, ES7$^s$ in yellow). **(B)** A close-up view of the *T.cruzi* ES7$^s$ and ES6$^s$ prior to (left) and post (right) eIF3 binding to the 40S **(C)** Overlay of mammalian and kinetoplastidian structures of individual eIF3 subunits with marked structural differences. The *T. cruzi* structures are depicted in dark and mammalian in light color shades. Curved arrows indicate the direction of *T. cruzi* eIF3 subunits structural rearrangement compared to their mammalian counterparts. Colored ovals highlight marked structural differences between *T. cruzi* and mammalian eIF3 subunits. **(D)** Cartoon representation of the eIF3 atomic model showing the eIF3 helical bundle in mammals (upper panel) and in *T. cruzi* (lower panel). Dark arrow indicates the shift of a helix from eIF3f in *T. cruzi* to compensate for the absence of eIF3m.

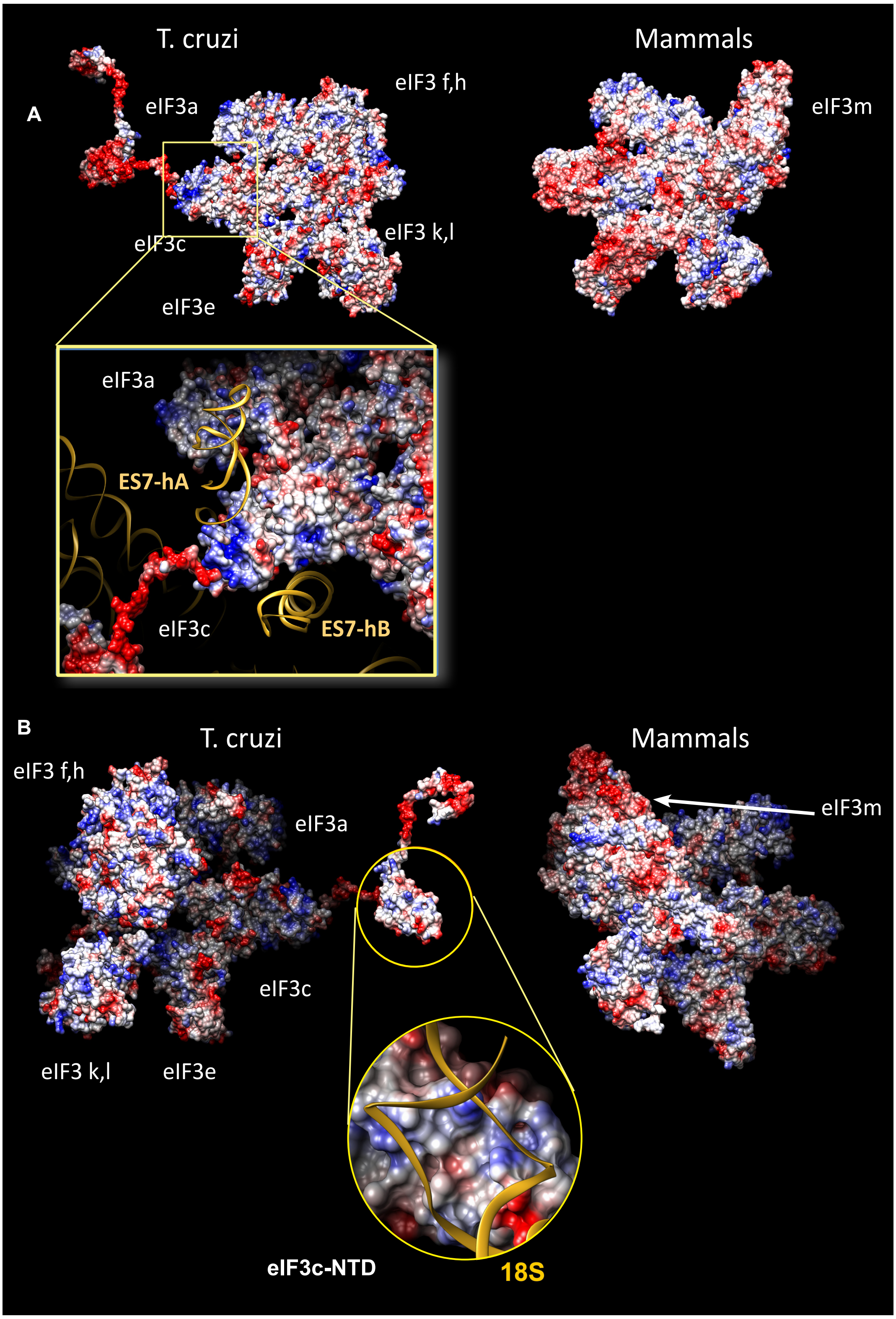

**Supplementary Figure 14. Charge surface analysis of the *T.cruzi* and mammalian eIF3 structures. (A)** Surface representation of the *T. cruzi* (left) and mammalian (right) eIF3 structure seen from the 40S platform side. Lower panel: close-up view of *T.cruzi* eIF3c and its interaction with the ES7$^s$ helix A and helix B. Model is color-coded according to the electrostatic potential – negative in red and positive in blue. **(B)** Surface representation of the *T. cruzi* (left) and mammalian (right) eIF3 structure seen from the 40S solvent side. Lower panel: close-up view of the *T.cruzi* eIF3c-NTD and its interaction with 18S RNA.

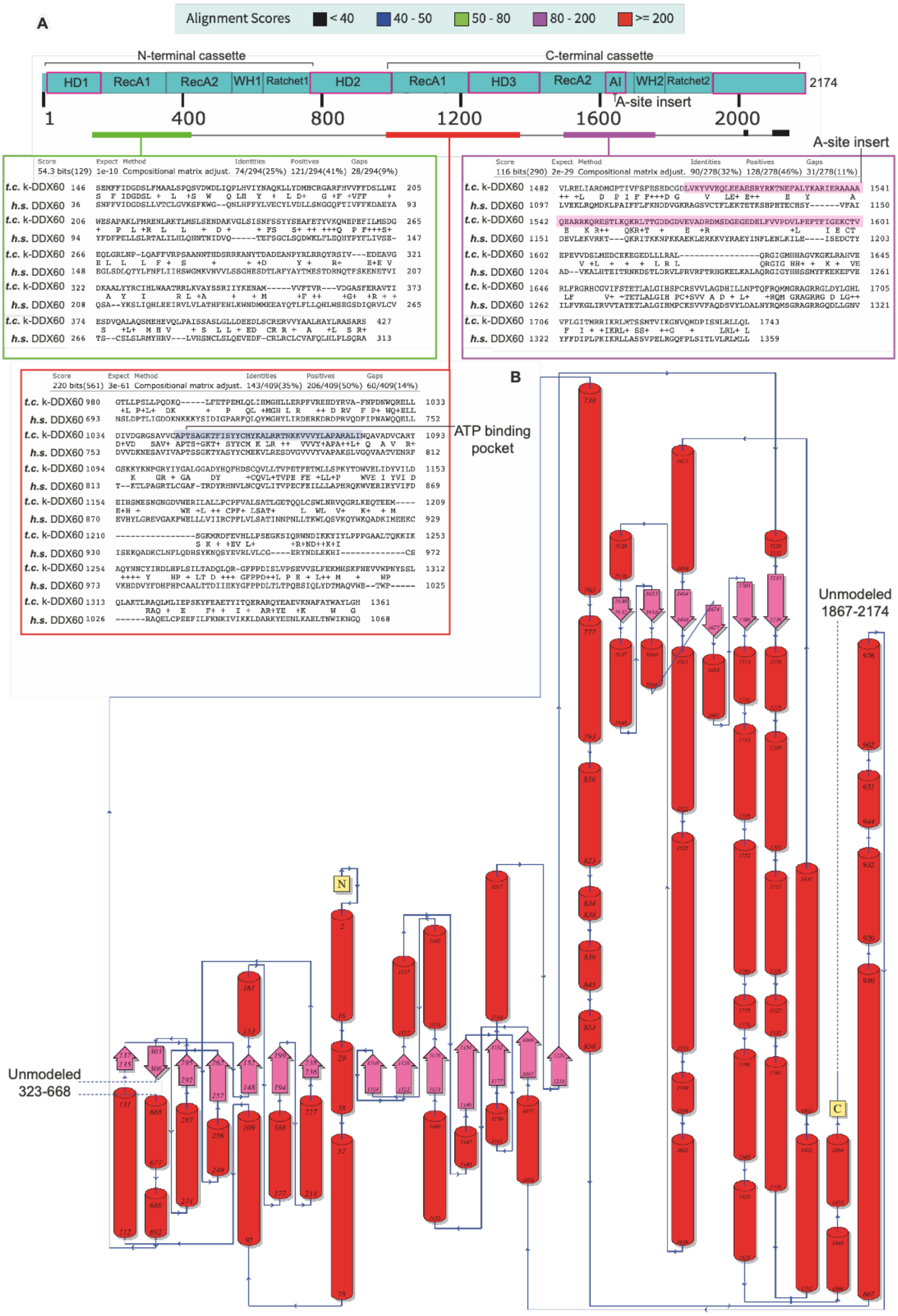

**Supplementary Figure 15. Sequence alignment between human DDX60 and k-DDX60, and its secondary structure.** (**A**) BlastP alignment between *T. cruzi* k-DDX60 and human DDX60 showing the relatively modest global homology between both proteins. Only most homologous regions were presented (in green, purple and red boxes). Magenta boxes on domains annotation schema highlight the trypanosomatid-specific domains that are inexistent in DDX60 from human and other eukaryotic species. Pink and violet colors highlight the A-site Insert (AI) and the ATP binding pocket in k-DDX60, respectively. (**B**) Secondary structure elements diagram for k-DDX60 based on its 3D model. Some parts couldn't be modeled.

| | Ribosomal RNA | Ribosomal protein | Initiation factors |
|---|---|---|---|
| **eIF1** | N65-G2303, C64-G2303, Q81-C2282, R33-A1341, R33-G2283, K37-G2283, R56-G2303, R61-C2183 | **none** | **eIF2-β : R29-S251, Q31-S327, Q43-T325, H27-T325, V77-Y326**<br>**eIF2-ɣ : S16-N459, V17-V147, Q12-Q412, L21-V85, Q13-V147**<br>**eIF3c : L49-F36, I54-W35 R53-E37, R52-T39, N96-R26, L49-I31** |
| **eIF1A** | N48-A2277, R66-C620, W74-A2279, **R155-G1685** | **eS30 :** E35-R10, F88-L8<br>**uS13 : D162-R119, L164-V124**<br>**uS19 : V158-V100, V158-A82, V158-A111**<br>**uS12 :** F88-L91 | **eIF2-β : Y133-L282, V134-N208, F135-P213, F135-Y279** |
| **eIF2-α** | **none** | **uS7 :**, Y200-K177, Y200-D180, Y200-R184, T148-R122, Y166-V120, T167-R122, D195-R184 | **tRNA :** K104-C55, R105-G52, R108-U54, W119-C55, H 232-C55, E296-U54<br>**eIF2-ɣ :** R331-E279, F315-L321, V320-L350, P350-F268 |
| **eIF2-β** | **R333-U1340, R333-G1342, R337-U1339, R337-U1340** | **uS19 :** N259-R137 | **tRNA :** K221-A36, N255-G25, K300-G68, R303-G69,<br>**eIF1 :** S251-R29, E267-Q32, T325-Q43, Y326-V77, Y326-H27, S323-R29<br>**eIF1A : N208-V134, P213-F135, Y279-F135, L282-Y133**<br>**eIF5 : N118-R265, L120-V329, L120-A262, L123-V325, K125-Q364, V132-W372, L142-F331**<br>**eIF2-ɣ :** N173-H248, T176-Y245, G181-Y241, Y182-Y211, Y184-D240, S185-N238, R189-E204, M305-E83, T317-M86, T317-E83 |
| **eIF2-ɣ** | **none** | **none** | **tRNA :** K79-C73, D269-A75, K272-A72, R282-A75<br>**eIF1 : V85-L21, I88-L21, V147-V17, N459-S16, Q412-Q12**<br>**eIF2-α :** E279-R331, L321- F315, L350- V320, F268- P350<br>**eIF2-β :** H248- N173, Y245- T176, Y241- G181, Y211- Y182, D240-Y184, N238- S185, E204- R189, E83- M305, M86- T317, E83-T317<br>**eIF5 : S224-R230, D219-R229, S220-R273, F383-L240, N430-T205, P431-D204, W465-T237, R469-T205**<br>**k-DDX60: P171-P770, D209-R902** |
| **eIF3c** | **S52-A1360, R53-C1361, K56-C1596, R127-C370, Q204-U1526, K207-A1525 R215-A1523, R232-U1476 and U1478, Q329-G1438, R331-U1439, R243-U1526** | **eS27 :** Q191-Q56, K192-E54 | **eIF1 : I31-M97, I31-L49, F36-L49, E37-R53, W35-F91, W35-I54, T39-R52**<br>**eIF3d : P234-W44, R295-W44, L489-W44, L233-A47, L380-F9, L418-W16, R419-P13, I434-M28, N437-D26**<br>**k-DDX60: N-ter tail with Y832 and F834** |
| **eIF3a** | | **eS1 :** T7-Q77, R8-T77, T12-R192, L17-I194 | |
| **eIF3d** | **D43-G1532, D50-A1475, R149-U1863 and U1862, R294-U1866 and C1867, D306-U1864,Q296-G1861, K301-U1863** | **eS27 : T36-K37, I39-F80, I39-L74**<br>**S33 :** R219-E76, D255-R83, K371-M98, Q368-K94, L435-M73<br>**uS7 :** Q434-E21, Q368-D26, E368-R51<br>**RACK1 :** S409-E277, N410-Q279 | **eIF3c : F9-L380, P13-R419, W16-L418, D26-N437, W44-P234, W44-R295, W44-L489, A47-L233, M28-I434**<br>**eIF3e : F3-T198, L5-A196, P6-T198, W16-I246, W16-Q247, E7-T245 ; P13-T248** |
| **eIF5** | **none** | **none** | **eIF2-β : A262-L120, R265-N118, V325-L123, V329-L120, I332-L142, Q364-K125, W372-V132**<br>**eIF2-ɣ : D204-P431, T205-R469, T205-N430, R229-D219, R230-S224, T237-W465, L240-F383, R273-S220**<br>**k-DDX60: D284-S944, D288-R941, K292-S826** |
| **k-DDX60** | **S26-U1722, R95-U1723, K724-A51, Q725-A51, H728-G477** | **eS12 : S3-D70, R6-E72**<br>**eS31 : Y5-K94, E92-L92, E93-K94**<br>**uS12 : R739-Q73, D744-N97 T740-N97** | **tRNA : Q1548-A34, S1551-A34**<br>**eIF2-ɣ : P770-P171, R902-D209**<br>**eIF3c : Y832 and F834 with N-ter tail**<br>**eIF5 : S826-K292, R941-D288, S944-D284** |

**Supplementary Table 1. Detailed overview of interactions between eIFs, ribosomal proteins, rRNA and k-DDX60.** Novel interactions revealed by analysis are colored in deep blue. Most of these novel interactions are shown in ribbons and sticks models fitted into their corresponding densities in Fig. S 7 and 8.

| Data Collection | *T. cruzi* 43S | *T. cruzi* 43S + ATP | *L. tarentolae* 43S |
|---|---|---|---|
| Microscope | Titan Krios | Titan Krios | Talos Artica |
| Voltage (kV) | 300 | 300 | 200 |
| Magnification | 127,272 | 127,272 | 120,000 |
| Pixel size (Å) | 1.1 | 1.1 | 1.21 |
| Detector | Gatan Summit K2 | Gatan Summit K2 | Falcon III |
| Defocus range (µm) | -0.6 to -4.5 | -0.6 to -4.5 | -0.6 to -3.0 |
| Tot. electron exposure ($e^{-}Å^{-2}$) | 30 | 30 | 40 |
| Exposure rate ($e^{-}Å^{-2}frame^{-1}$) | 1.5 | 1.5 | 2.0 |
| Data collection software | SerialEM | EPU | EPU |
| **Data Processing** | | | |
| Independent data collections | 1 | 1 | 1 |
| Useable micrographs | 3271 | 2638 | ? |
| Particles | 202920 | 98840 | 52302 |
| Final particles (43S) | 33775 | 19700 | 10144 |
| Accuracy translations (pix) / rotations (°) | 0.432/0.234375° | 0.432/0.234375° | 0.432/0.234375° |
| Resolution (Å, 0.143 FSC) | 3.33 | 4.3 | 8.1 |
| Local resolution range (Å) | 2.5-6 | N/A | N/A |
| **Model Composition** | | | |
| Chains | 59 | N/A | N/A |
| Non-hydrogen atoms | 136893 | N/A | N/A |
| Protein residues | 11196 | N/A | N/A |
| RNA bases | 2225 | N/A | N/A |
| **Refinement** | | | |
| Software | Phenix_ValidationEM | N/A | N/A |
| Resolution (Å) | 3.33 | N/A | N/A |
| CC (mask) | 0.57 | N/A | N/A |
| CC (main chain) | 0.5 | N/A | N/A |
| CC (side chain) | 0.58 | N/A | N/A |
| **R.M.S deviations** | | | |
| Bond lengths (Å) | 0.020 | N/A | N/A |
| Bond angles (°) | 2.209 | N/A | N/A |
| **Validation** | | | |
| Molprobity score | 2.65 | N/A | N/A |
| Clashscore, all atoms | 9.20 | N/A | N/A |
| Rotamers outliers (%) | 3.48 | N/A | N/A |
| Cβ outliers (%) | 1.41 | N/A | N/A |
| CaBLAM outliers (%) | 10.9 | N/A | N/A |
| **Ramachandran plot** | | | |
| Favored (%) | 79.09 | N/A | N/A |
| Allowed (%) | 14.19 | N/A | N/A |
| Outliers (%) | 6.72 | N/A | N/A |

**Supplementary Table 2: Data collection, processing, refinement and model statistics.** A near complete atomic model was only derived for the highest resolution cryo-EM reconstruction, i.e. the *T. cruzi* 43S PIC stalled with GMP-PNP. The *T. cruzi* 43S PIC stalled with GMP-PNP supplemented with ATP presents a lower resolution and therefore we didn't derive a full atomic model, instead we flexibly fitted k-DDX60 only into its map to illustrate its conformational changes.
